# A lncRNA drives developmentally-timed decay of all members of an essential microRNA family

**DOI:** 10.1101/2025.07.30.667716

**Authors:** Acadia L. Grimme, Lu Li, Alyshia Scholl, Bridget F. Donnelly, Neha Channamraju, Karl-Frédéric Vieux, Lecong Zhou, Geraldine Seydoux, Mingyi Xie, Katherine McJunkin

## Abstract

The spatiotemporal expression patterns of microRNAs (miRNAs) are crucial to their function. Target-directed miRNA degradation (TDMD) is an emerging regulatory module that contributes to these expression patterns wherein a specialized RNA (TDMD trigger) drives miRNA decay through base pairing and resulting recruitment of E3 ubiquitin ligase ZSWIM8/EBAX-1. Extensive base pairing to the miRNA seed region and 3’ end has been proposed as a key feature that distinguishes TDMD triggers from conventional mRNA targets of miRNAs, which primarily pair with the seed. Here we identify the long noncoding RNA, *tts-2*, as a TDMD trigger for *mir-35-42*, the most abundant miRNA family in *C. elegans* early embryos. We demonstrate that a single site in *tts-2* drives decay through base pairing with the seed sequence shared by all eight family members. A second site in *tts-2* supports decay of *mir-38* with incomplete seed complementarity. Our findings demonstrate that extended base pairing is not a universal requirement for TDMD, and that TDMD drives developmentally-timed clearance of abundant miRNAs at the exit of *C. elegans* embryogenesis.

## Introduction

microRNAs (miRNAs) are small noncoding RNAs which post-transcriptionally repress mRNAs through partial base pairing to sites in the mRNAs’ 3′ UTRs^1^. To carry out this function, each miRNA is bound to an Argonaute (Ago) protein to form the core of a <u>miR</u>NA-induced <u>s</u>ilencing <u>c</u>omplex (miRISC)^2^. Because miRNAs function in a dose-dependent manner, their abundance is tightly controlled through regulated transcription, biogenesis, and decay^3^. Despite the crucial role of miRNAs in regulating gene expression, the understanding of their regulated decay mechanisms has lagged far behind that of their transcription and biogenesis.

Individual miRNAs display complex spatio-temporal expression patterns, and accordingly, their developmental stage- and cell type-specific expression drives or reinforces stage- and cell type-specific protein-coding gene output^4^. Beyond just its contribution to tuning steady state miRNA levels, regulated miRNA decay must also contribute to rapid shifts in miRNA repertoires across the time scales of development and differentiation.

This is especially true given the fact that miRNAs are generally long-lived species (average half-lives > 24h) due to their protective physical interactions with Ago proteins which shield the 5′ phosphate and 3’ hydroxyl of the miRNA in binding pockets in the MID and PAZ domains, respectively^5^. Despite their generally long half-lives, miRNAs also display a broad sequence- and context-dependent variance in decay rates^6–15^. To date, most known instances of sequence-dependent miRNA decay are driven by target-<u>d</u>irected <u>m</u>iRNA <u>d</u>egradation (TDMD)^16^.

TDMD is elicited by the interaction of the miRNA with specialized targets, often called TDMD “triggers” to distinguish them from conventional mRNA targets^17^. Interactions between miRNAs and conventional targets are largely driven by base pairing between the target and the “seed” region of the miRNA (positions 2-8 from the 5′ end), with a frequently-observed supportive role of base pairing between the target and miRNA positions 13-16 (the “supplemental” region)^1^. Known TDMD triggers base pair more extensively with the miRNA that they degrade, especially in the 3′ extremity of the miRNA ^18–28^. A prevailing view in the field is that this extensive base pairing is what distinguishes the outcome of a miRNA interaction with a TDMD trigger (miRNA decay) from that with a conventional target (target repression and decay)^29^.

Interactions between miRISC and known TDMD triggers has been proposed to result in three outcomes. 1) Extensive base pairing with the trigger pulls the miRNA 3′ end out of the Ago PAZ domain; this conformational shift in the miRNA propagates to a larger overall shift in the Ago protein conformation, allowing its bilobed structure to open, broadening the central cleft^30^. 2) The release of the miRNA 3’ end from the PAZ domain exposes it to remodeling enzymes: terminal nucleotidyl transferases (TNTs) that add untemplated terminal nucleotides (tailing) and exonucleases that trim off one or more terminal nucleotides^20^. This is collectively termed target-<u>d</u>irected tailing and trimming (TDTT). 3) An E3 ubiquitin ligase containing the substrate recognition module ZSWIM8 (<u>Z</u>inc finger <u>SWIM</u>-type containing <u>8</u>) is recruited to the miRISC, driving ubiquitination of the Ago and proteasome-dependent decay of the miRNA (presumably via Ago decay and total exposure of the miRNA to cellular nucleases)^31,32^. While TDTT is genetically and biochemically separable from TDMD^27,31,32^, most models currently propose that the Ago conformational change drives the recruitment of ZSWIM8 and resulting TDMD.

In this work, we elucidate the developmentally timed decay of a group of essential miRNAs. Because of their shared seed sequence and resulting redundant function on a set of common targets, these miRNAs, *mirs-35-42* are termed the *mir-35-42* family. These miRNAs are the most abundantly expressed miRNAs in early *C. elegans* embryos, and knockout of all family members results in embryonic arrest and lethality (or a host of milder phenotypes if a subset of the family members is deleted)^33–42^. All eight miRNAs are sharply degraded in late embryogenesis, suggesting their decay is accelerated by a sequence-specific process^41,43^. We previously showed that the decay of these miRNAs depends on their seed sequence, but not on other portions of the sequence outside this region^44^. Surprisingly, given the limited sequence-dependence, *mir-35-42* decay at the end of embryogenesis also depends on the ortholog of TDMD factor ZSWIM8, known as EBAX-1 in *C. elegans* ^32,44–46^.

Here we show that the decay of *mir-35-42* is driven by EBAX-1-dependent TDMD. The TDMD trigger that drives decay of all eight family members is a lncRNA, *tts-2*, whose expression peaks in late embryogenesis. Two binding sites in *tts-2* are responsible for degrading all eight family members, with one site being wholly responsible for the decay of seven family members and both sites acting redundantly to degrade the eighth family member. We extensively mutate the primary *tts-2* binding site, demonstrating that this TDMD is dependently only on base pairing to the miRNA seed region. This work demonstrates that extensive base pairing with the miRNA – and the resulting conformation shift in Ago – is not a necessary feature that distinguishes TDMD triggers. Instead, supplemental or alternative cues must mark some (if not all) trigger-miRISC ternary complexes for E3 recruitment and TDMD.

## Results

### EBAX-1 is required for seed-mediated decay of *mir-35-42*

The decay of *mir-35* at the end of embryogenesis depends on 1) the *mir-35* seed region alone and 2) the ZSWIM8 ortholog EBAX-1. To determine whether these two requirements reflect the same or different decay pathways, we performed a series of small RNA-seq (sRNA-seq) experiments to determine the sensitivity of wild type and mutant variants of *mir-35* to loss of EBAX-1. We first confirmed that the loss of EBAX-1 stabilizes *mir-35-42* at the transition from embryo to the first larval stage, L1 (EtoL1), as previously reported (Figure 1A, Tables S5-S6)^14,44,46^. *mir-35-42* represent a vast majority of miRNAs stabilized in *ebax-1(null)* L1s; while eight out of thirteen EBAX-1-sensitive miRNAs in L1s are *mir-35-42*, they account for 88% of differentially stabilized miRNA reads due to their high abundance (Figure 1B-C, Table S6)^46^.

**Figure 1.**
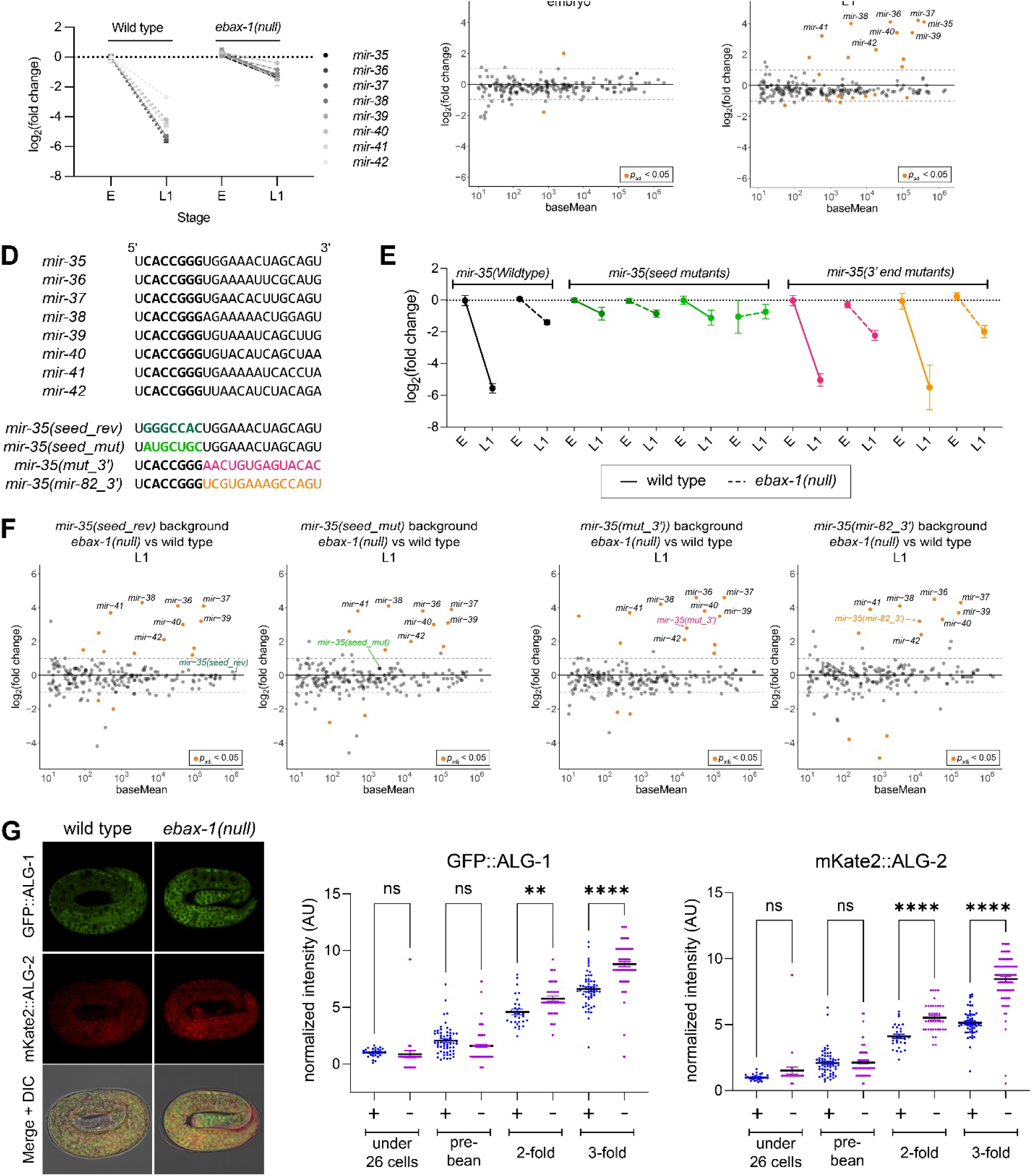
The *mir-35* seed sequence is sufficient for degradation by EBAX-1. (A) miRNA levels (sRNA-seq) normalized to embryonic wild type levels. E, embryo. Mean and SEM of three biological replicates shown. (B and C) MA plots of sRNA-seq from *ebax-1(null)* vs wild type animals at the indicated developmental stages. (D) Sequences of *mir-35-42* and the *mir-35* sequence variants used in this study. (E) miRNA levels (sRNA-seq) normalized to the embryonic levels of the corresponding *mir-35* sequence. Mean and SEM of three biological replicates shown. (F) MA plots of the L1 sRNA-seq in *ebax-1(null)* vs wild type in strains containing the indicated *mir-35* variant allele. (G) Left: Representative images of GFP::ALG-1 and mKate2::ALG-2 in 3-fold embryos in wild type or *ebax-1(null)* background. Right: Quantification of fluorescent signal from GFP::ALG-1 or mKate2::ALG-2 across different stages of embryonic development. One way ANOVA with Sidak’s multiple comparison test. **, p=0.0099; ****, p<0.0001. (A-F) sRNA-seq was normalized using total genome-mapping reads. sRNAseq analysis was carried out with 3 biological replicates per genotype, and miRNAs with baseMean ≥ 10 reads are shown.

We previously showed that two variants of *mir-35* in which the seed sequence is mutated (at the endogenous genomic locus) are not degraded at EtoL1 (Figure 1A)^44^. When these *mir-35* seed mutations were combined with *ebax-1(null)*, the attenuation of decay was similar to that observed in the context of either single mutant (*ebax-1(null)* or *mir-35* mutation) (Figure 1D-F). This observation suggests that *mir-35* seed mutations and *ebax-1(null)* disrupt the same decay pathway.

Next, we tested a set of *mir-35* variants in which all nucleotides 3′ of the seed sequence were mutated. One of these (*mut_3′*) was designed to differ from wild type *mir-35-42* while preserving overall GC content (Figure 1D). The other (*mir-35(mir-82_3′)*) swaps in the corresponding region of another miRNA expressed in embryos that does not undergo EtoL1 decay (*mir-82*) (Figure 1D). We previously showed that these sequence changes do not disrupt EtoL1 decay (though they do result in higher embryo levels, which are not shown here due to normalization required by use of sRNA-seq rather than absolute quantification qPCR)^44^. When combined with the *ebax-1(null)* mutation, the 3′ end variants are strongly stabilized at the EtoL1 (Figure 1E-F). Because these variants are mutated in all positions 3′ of the seed sequence, we conclude that the *mir-35-42* seed sequence is sufficient for EBAX-1-dependent *mir-35-42* decay at the EtoL1 transition.

### Both ALG-1 and ALG-2 are sensitive to EBAX-1

Given that EBAX-1 orthologs can selectively target Ago paralogs ^47^, we tested whether the identity of the Ago protein may play a role in this degradation process. *C. elegans* contains two paralogous proteins that are responsible for the vast majority of miRNA loading, <u>A</u>rgonaute-like <u>g</u>ene 1 (ALG-1) and ALG-2^48–52^. *mir-35-42* are predominantly loaded in ALG-2^42,53,54^. Seed mutant variants of *mir-35* are also preferentially loaded in ALG-2, suggesting that altered loading does not underlie their perdurance at EtoL1 and demonstrating that ALG-2 loading is not sufficient for EtoL1 decay (Figure S1A).

To examine whether ALG-2 loading is required for *mir-35-42* decay, we examined knockout mutants of ALG-2 and ALG-1 using sRNA-seq. As expected, many miRNAs were present at lower levels in both mutants (Figure S1B, Tables S7-S10). In the ALG-2 knockout, levels of *mir-35-42* family members were decreased, as previously observed^38^ (Figure S1B, Tables S9-S10). However, when examining the extent of decay at the EtoL1 transition, the amplitude of decay was remarkably similar to wild type in *alg-2(null)* (Figure S1C). This indicates that specific loading into ALG-2 is not required for EtoL1 decay. In the ALG-1 knockout, *mir-35-42* levels and EtoL1 decay were very similar to wild type (reflecting their predominant loading in ALG-2) (Figure S1D-E, Tables S7-S8).

Overall stabilization of Ago has not yet been observed in ZSWIM8 mutants in other species, presumably due to the small portion of the miRISC pool that contains ZSWIM8-sensitive miRNAs in each context. We reasoned that because *mir-35-42* represent such a major proportion of all miRNAs in *C. elegans* embryos, this context may allow for changes in overall levels of ALG-1/2 in the absence of EBAX-1. To determine whether EBAX-1-dependent decay affects the levels of ALG-1 and ALG-2, we quantified levels of ALG-1/2 tagged with fluorescent proteins knocked in at their endogenous genomic loci^54^. We first confirmed that the N-terminal tagged Ago alleles do not disrupt *mir-35-36* decay at the EtoL1 transition (Figure S1F). We then measured tagged ALG-1 and ALG-2 levels across embryonic development by imaging. While no difference was observed in early embryos, ALG-1 and ALG-2 levels were 33% and 64% higher, respectively, in *ebax-1(null)* compared to wild type late-stage embryos, coincident with the onset of *mir-35-42* decay (Figure 1G). Increased levels were observed ubiquitously across many tissues, and the amplitude of ALG-2’s increase was greater than that of ALG-1 (Figure 1G). Since *mir-35-42* account for 87% or 88% of differentially stabilized miRNA reads in *ebax-1(null)* late-stage embryos and L1s^44,46^, respectively, these observations suggest that *mir-35-42* ALG-1/2 miRISCs undergo a decay process analogous to known instances of TDMD, proceeding via EBAX-1-mediated Ago degradation.

### Chimeric eCLIP identifies *tts-2* as a candidate *mir-35-42* TDMD trigger RNA

To identify the TDMD trigger RNA driving *mir-35-42* decay, we used chimeric <u>e</u>nhanced <u>c</u>rosslinking immuno<u>p</u>recipitation (chimeric eCLIP^55^), a method similar to <u>c</u>rosslinking <u>a</u>nd <u>s</u>equencing of <u>h</u>ybrids (CLASH)^56^, which has previously successfully identified TDMD triggers^26,28,57^. This workflow favors intermolecular ligation of miRNA-RNA fragments, resulting in hybrid reads, allowing for identification of the paired molecules (Figure 2A). We generated samples for chimeric eCLIP using a strain in which ALG-2 is 3xFLAG-tagged at its genomic locus^58^. Mixed embryo samples were used since the onset of *mir-35-42* decay occurs in late embryogenesis. We further reasoned that loss of EBAX-1 should result in stabilization of the miRISC-TDMD trigger RNA interaction due to the loss of Ago degradation central to TDMD; we therefore generated samples in wild type and *ebax-1(null)* backgrounds (Figure 2A).

**Figure 2.**
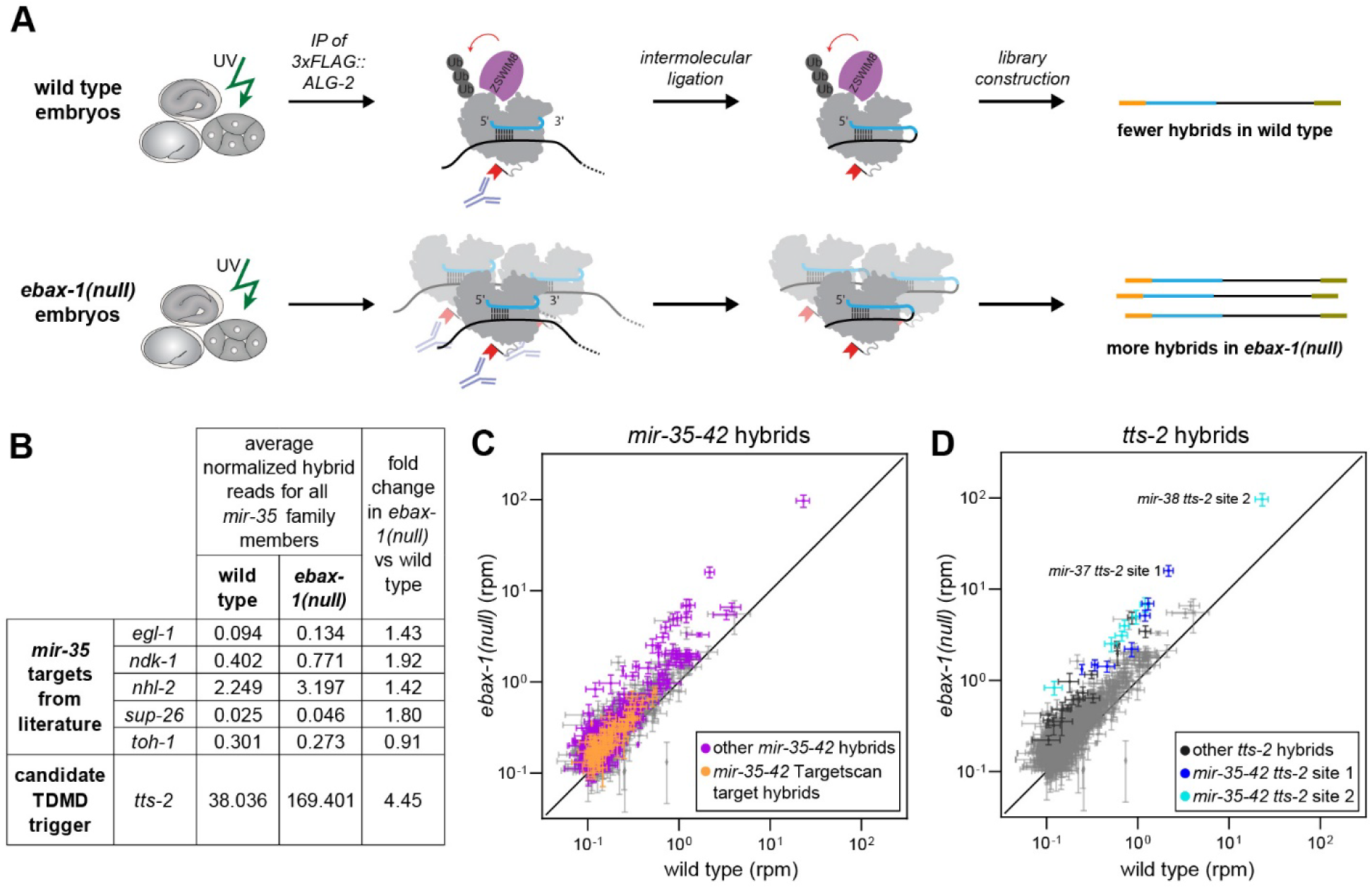
Long-noncoding RNA *tts-2* identified as a candidate *mir-35* family TDMD trigger. (A) Schematic depicting chimeric eCLIP in wild type and *ebax-1(null)* embryos. (B) Normalized average hybrid reads (reads per million, RPM) across four biological replicates per genotype for known *mir-35* targets and *tts-2*. (C-D) Average hybrid reads (RPM) highlighting all *mir-35-42* hybrids (C) or all *tts-2* hybrids (D). Mean RPM +/-SEM are shown.

mRNAs and non-ribosomal noncoding RNAs accounted for 42% and 49% of hybrid reads in wild type and *ebax-1(null)* libraries, respectively (Figure S2). Both conventional targets and TDMD triggers are expected to be preferentially isolated in *ebax-1(null)* samples due to increased abundance of *mir-35-42*. Indeed, previously-studied targets and predicted targets of *mir-35-42* displayed a modest increase in hybrid reads with *mir-35-42* in the *ebax-1(null)* samples compared to wild type (Figure 2B-C). By examining abundant hybrid reads (>0.1 reads per million) with the greatest enrichment in *ebax-1(null)* samples compared to wild type, a candidate *mir-35-42* TDMD trigger was immediately apparent. Among the top 25 most enriched hybrids in *ebax-1(null)* relative to wild type, nineteen were hybrids of *mir-35-42* and the lncRNA *transcribed telomerase <u>s</u>equence (tts-2)* (Table S11). In particular two binding sites in *tts-2* represented multiple of the most abundant and most *ebax-1(null)*-enriched hybrids (Figure 2B,D). Both of these sites in *tts-2* were cloned in hybrids with all eight of the *mir-35-42* family members and are conserved in related nematode species (Figure 3A), and abundant *tts-2/mir-35-42* hybrids were together enriched 4.45-fold in *ebax-1(null)* compared to wild type.

**Figure 3.**
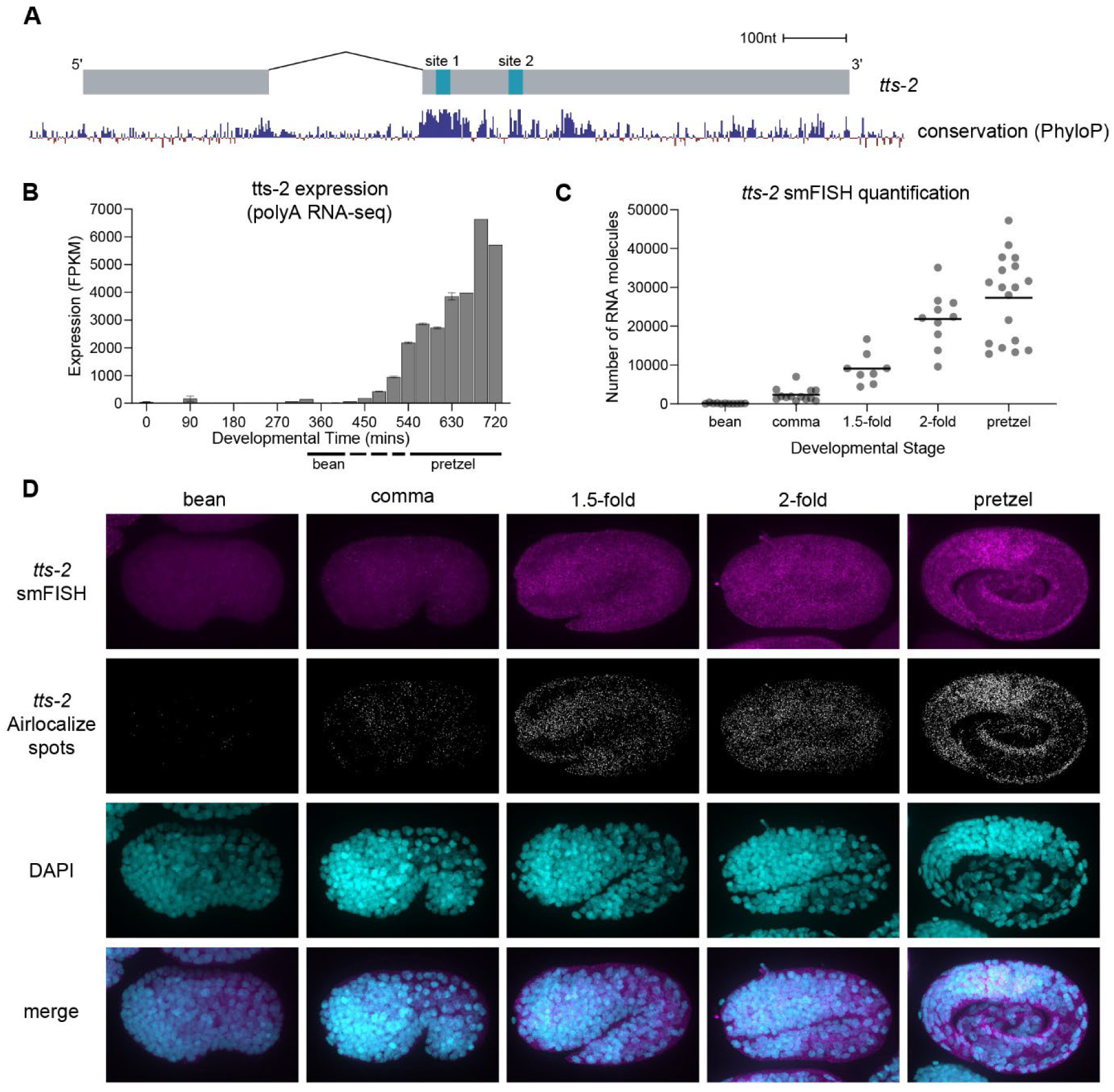
*tts-2* expression peaks in late embryogenesis. (A) *tts-2* gene model with PhyloP conservation track from UCSC genome browser. (B) Expression of *tts-2* across embryonic development from polyA RNAseq data (modENCODE), shown as mean +/-SEM (Gerstein et al. 2010). (C) Quantification of *tts-2* smFISH signal by Airlocalize across embryonic development. Individual embryo values and mean shown. (D) Representative images of *tts-2* smFISH and Airlocalize spots.

*tts-2* is annotated in WormBase as a mature spliced transcript of 882 nt^59^. *tts-2* is readily captured in high abundance in poly(A)-dependent RNA-seq datasets, and like most *C. elegans* Pol II transcripts, is transpliced at its 5′ end to the splice leader 1, SL1 (Figure S3). Functional analysis of *tts-2* has not been reported.

### *tts-2* expression peaks in late embryogenesis

If *tts-2* expression determines the timing of *mir-35-42* decay, then we would expect it to be lowly expressed in early embryogenesis and highly expressed in late embryogenesis. Published RNA-seq data in *C. elegans* embryos shows that *tts-2* displays exactly such an expression pattern (Figure 3B)^60^. To confirm this expression pattern and examine the tissues of expression, we performed single molecule FISH (smFISH). Specificity of the *tts-2* FISH signal was confirmed by its absence in a *tts-2(null)* mutant (Figure S4). We observed that *tts-2* is lowly expressed in early embryos (up through the bean stage) but rapidly increases in expression in late embryos, with the average expression in pretzel-stage animals 37.6-fold higher than at bean stage (Figure 3C-D). *tts-2* was observed in the cytoplasm and broadly across many if not all tissues (Figure 3D). A GFP reporter knocked in at the 5′ end of *tts-2* shows a similar pattern of expression, ramping up sharply in late embryogenesis (Figure S5A-B). We conclude that *tts-2* is expressed at the right time and place during embryogenesis to function as a *mir-35-42* TDMD trigger.

### *tts-2* drives decay of *mir-35-42*

To test whether *tts-2* is required for *mir-35-42* decay, we generated a 1485-bp knockout of the *tts-2* gene [*tts-2(null)*] and measured *mir-35-42* levels by sRNA-seq in L1s. Loss of *tts-2* stabilized all eight members of the *mir-35-42* family in L1s (Figure 4A-B, Table S12). The *mir-35-42* passenger strands (byproducts of *mir-35-42* biogenesis also captured in sequencing) were not upregulated, indicating that *tts-2* acts on *mir-35-42* at the level of decay, since transcription- or biogenesis-level effects would be expected to impact both strands (Figure 4A). No other miRNAs are impacted by *tts-2(null)* in L1s (Figure 4B).

**Figure 4.**
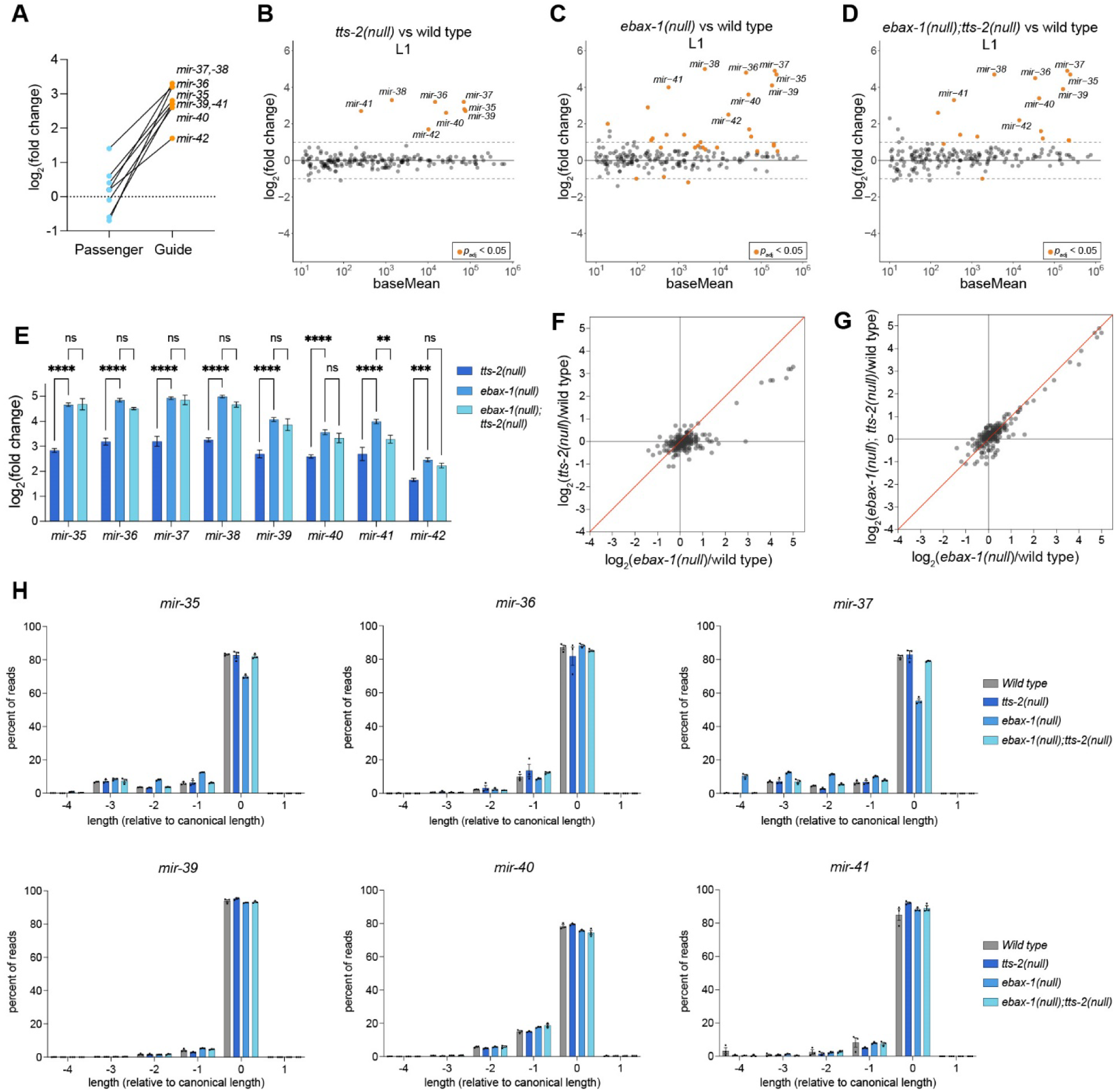
*tts-2* acts with EBAX-1 to drive *mir-35-42* decay. (A) Log_2_(fold change) of *mir-35-42* passenger and guide strands in *tts-2(null)* versus wild type (sRNA-seq). (B-D) MA plots of sRNA-seq for the indicated genotypes in L1 stage animals. (E) Comparison of log_2_(fold change) values in L1s for indicated genotype versus wild type. Mean +/- SEM shown. Two-way ANOVA with Dunnett’s test, **, p<0.01, ***, p<0.001, ****, p<0.0001. (F and G) Comparison of all log_2_(fold change) values for indicated genotypes relative to wild type. (H) Analysis of miRNA trimming. (A-G) sRNA-seq was performed with three biological replicates per genotype and normalized using total piRNA reads. miRNAs with baseMean ≥ 10 are shown.

The involvement of both EBAX-1 and *tts-2* suggests that *mir-35-42* decay is EBAX-1-dependent TDMD. However, the amplitude of *mir-35-42* stabilization in *tts-2(null)* is slightly lower than that observed in *ebax-1(null)* (Figure 4A-C,E-F, Tables S12-S13). This difference in potency could be due to a) the existence of secondary TDMD triggers acting with EBAX-1 to drive *mir-35-42* decay or b) two distinct decay mechanisms involving *tts-2* and *ebax-1*, respectively. To confirm that EBAX-1 and *tts-2* act through a common pathway, we compared the effects of each single mutant to *tts-2(null); ebax-1(null)* double mutant L1s. If *tts-2* functions as a trigger RNA in EBAX-1-dependent TDMD, then a *tts-2(null); ebax-1(null)* mutant should phenocopy the effect size observed in the *ebax-1(null)* background, whereas two independent degradation mechanisms would likely have additive effects on *mir-35-42* decay. The *tts-2(null); ebax-1(null)* sRNAseq results closely phenocopy the changes we see in the *ebax-1(null)* dataset, indicating that *tts-2* functions in the same pathway as EBAX-1 to drive *mir-35-42* degradation (Figure 4C-E, G, Table S14). We conclude that EBAX-1-dependent decay of *mir-35-42* depends primarily on *tts-2*, and likely also on secondary, less potent triggers.

### *tts-2* induces neither tailing nor trimming of most *mir-35-42* family members

TDMD triggers that extensively base pair with the miRNA 3’ end drive a conformational change that generally results in target-directed tailing and trimming (TDTT) due to loss of association between the miRNA 3’ end and the Ago PAZ domain^19,20,27,30,61,62^. Although TDTT is not required for TDMD, these modifications report on the exposure of the miRNA 3′ end and the concomitant Ago conformation.

We analyzed deep sequencing data in *ebax-1(null), tts-2(null)* and double *ebax-1(null); tts-2(null)* mutants to determine whether TDTT is also observed in the case of *mir-35-42* TDMD. We detected no significant changes in tailing among any of the genotypes tested (Figure S6A). We observed increased trimming for *mir-35* and *mir-37* in *ebax-1(null)* (Figure 4H), consistent with a recent report from Stubna, et al^46^. Interestingly, the increased trimming was not observed in the *tts-2(null)* single mutant nor in the *ebax-1(null); tts-2(null)* double mutant (Figure 4H). Thus, the increased trimming of *mir-35* and *mir-37* observed in *ebax-1(null)* is driven by *tts-2*.

Notably, we did not observe changes in trimming for the other *mir-35-42* family members in *ebax-1(null), tts-2(null)* or the *ebax-1(null); tts-2(null)* double mutant (Figure 4H, Figure S6B). To determine whether the difference in TDTT could reflect differences in base pairing architecture with *tts-2*, we examined the binding sites in *tts-2*. Indeed, the binding site containing a perfect seed match to *mir-35-42* (“site 1”) has extensive predicted base pairing to the 3′ ends of *mir-35* and *mir-37*, but not the other *mir-35-42* family members (Figure 5A). Thus, *tts-2* TDTT correlates with the extent of miRNA 3′ end pairing to site 1, and for most *mir-35-42* family members, *tts-2*-driven TDMD occurs in the absence of TDTT.

**Figure 5.**
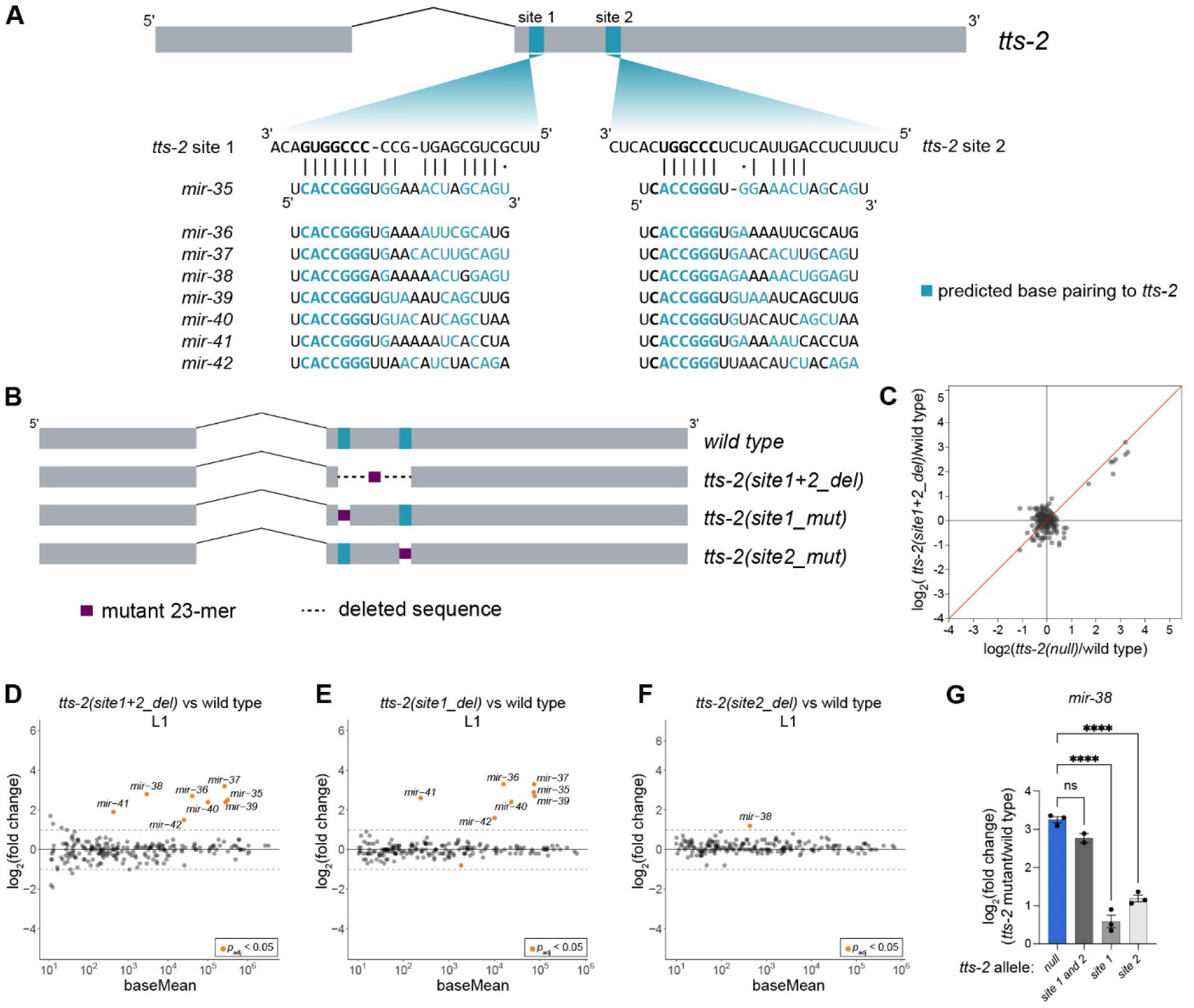
One binding site in *tts-2* is primarily responsible for *mir-35-42* degradation. (A) Predicted base pairing between *mir-35-42* and *tts-2*. (B) CRISPR generated mutations of each binding site in *tts-2*. (C) Comparison of all log_2_(fold change) values for indicated genotypes relative to wild type. (D-F) MA plots of sRNA-seq for the indicated *tts-2* mutants in L1 animals. (G) Log_2_(fold change) of *mir-38* in *tts-2(null)* and single binding site mutants compared to wild type. One-way ANOVA with Dunnett’s test; ****, p<0.0001. (C-G) sRNA-seq was performed with three biological replicates per genotype, except for *tts-2(site1+2_del)* (two replicates), and normalized using total piRNA reads. miRNAs with baseMean ≥ 10 are shown.

### *mir-35-42* decay is primarily driven by one binding site in *tts-2*

To define the features of *tts-2* required for TDMD, we first examined the binding sites for *mir-35-42* identified by chimeric eCLIP. As mentioned above, the first site (“site 1”) contains a perfect complement to the *mir-35-42* seed sequence and variable amounts of predicted base pairing to the 3′ ends of the family members (Figure 5A). The second site (“site 2”) has a near-complement to the *mir-35-42* seed, with predicted binding to miRNA positions 3-8. Pairing of the 3′ portion of *mir-38* to site 2 is extensive, whereas 3′ pairing of other family members to this site is minor (Figure 5A).

We first deleted both sites and the intervening region to determine if these sites are responsible for *mir-35-42* TDMD. sRNA-seq in L1s showed that this mutation (site1+2_del) resulted in stabilization of all eight family members (Figure 5B,D, Table S15). The amplitude of stabilization was similar to that observed in *tts-2(null)*, demonstrating that these sites are strictly required for *tts-2* function in TDMD (Figure 5C).

Next, we examined the contribution of each site individually (Figure 5B). Mutation of site 1 resulted in stabilization of all *mir-35-42* family members except *mir-38*, with amplitude similar to *tts-2(null)* and *tts-2(site1+2_del)* (Figure 5E, Table S16). Mutation of site 2 resulted in stabilization of *mir-38* alone (Figure 5F, Table S17). However, *mir-38* stabilization in the site 2 mutant was very mild compared to that observed in *tts-2(null)* and *tts-2(site1+2_del)* (Figure 5G). Therefore, site 1 and site 2 act redundantly to induce *mir-38* decay, since both sites must be mutated to observe robust stabilization of *mir-38*.

Among known examples of miRNA-TDMD trigger pairs, most trigger sites act solely in TDMD, whereas others also simultaneously serve as a site for miRNA-mediated target repression^23,27,31^. To determine whether *tts-2* is targeted by *mir-35-42,* we quantified levels of *tts-2* by qPCR in wild type, *ebax-1(null),* and the *tts-2* binding site mutants. We found no significant change in abundance when compared to wild type, indicating both that *tts-2* is not destabilized by binding of *mir-35-42* and that loss of TDMD in *tts-2* binding site mutants is not due to loss of expression (Figure S8A).

### *tts-2* is a highly potent TDMD trigger

Among known TDMD trigger interactions, a wide range in efficiencies has been observed. Whereas many TDMD triggers appear to degrade approximately a single miRNA, the lncRNA Cyrano/Oip5-as1 degrades ∼17 miRNA molecules per molecule of Cyrano^23,27,62,63^. To determine the relative stoichiometry of *tts-2* and *mir-35-42*, we performed qPCR using in vitro transcribed (*tts-2*) or synthetic RNA (*mir-35*) as standards for absolute quantification (Figure S7). Since our system is not steady-state, estimates of the ratio of *tts-2* to *mir-35-42* are approximate. When using L1 samples to estimate the number of molecules of *mir-35-42* degraded per molecule of *tts-2*, we observe that wild type L1s contain ∼192,252 copies of *mir-35* per ng of total RNA. Extrapolating based on known ratios of *mir-35-42* family members and fold-change observed in *tts-2(null)*, we estimate that approximately 6.39M more molecules of *mir-35-42* are present in 1ng of *tts-2(null)* L1 RNA compared to wild type. Given the measurement of ∼89,836 molecules of *tts-2* per ng of total L1 RNA, we estimate that each copy of *tts-2* is responsible for degrading ∼71 molecules of *mir-35-42*. Since *tts-*2 peaks earlier in development, we also used absolute measurements of *tts-2* in mixed embryo samples fitted to the distribution of stages present in those samples and the temporal profiles of *tts-2* to estimate the peak of *tts-2* (∼346,742-398,941 molecules per 1ng of late-stage embryo RNA). Using this estimate, the ratio remains high: 16-18.4 molecules of *mir-35-42* degraded per molecule of *tts-2*, making *tts-2* comparable to the strongest TDMD trigger reported to date.

### Seed pairing to *tts-2* is sufficient for TDMD

To further examine the sequence requirements for *tts-2* driven TDMD, we made point mutations across binding site 1 in *tts-2.* Ten mutant strains were generated in which various subsets of nucleotides in *tts-2* were mutated, spanning the position opposite miRNA nucleotide 8 (t8) to 27 bp further upstream in *tts-2* (t35) (Figure 6A). Four of these mutants were assayed by deep sequencing. Mutations outside the region predicted to base pair with the miRNAs (t26-35) had no effect on *mir-35-42* decay (Figure 6A-B, Table S18). Of the mutations in the miRNA base pairing region, only the mutation overlapping with the seed pairing region (t8-13) had dramatic effects on *tts-2* function, mimicking the site 1 deletion mutant (Figure 6C, Table S19). Mutation at t14-19 showed very mild stabilizing effects for *mir-36* and *mir-37* (Figure 6D, Table S20). However, broader mutations encompassing this region (t9-20) had no effect on decay, despite their rational design to maximally disrupt base-pairing to *mir-35-42* (Figure 6E, Table S21). Thus, we speculate that the t14-19 mutation is idiosyncratic in some way, perhaps introducing structures that impede miRNA binding since qPCR showed wild type-like *tts-2(t14-19)* expression levels (Figure S8A).

**Figure 6.**
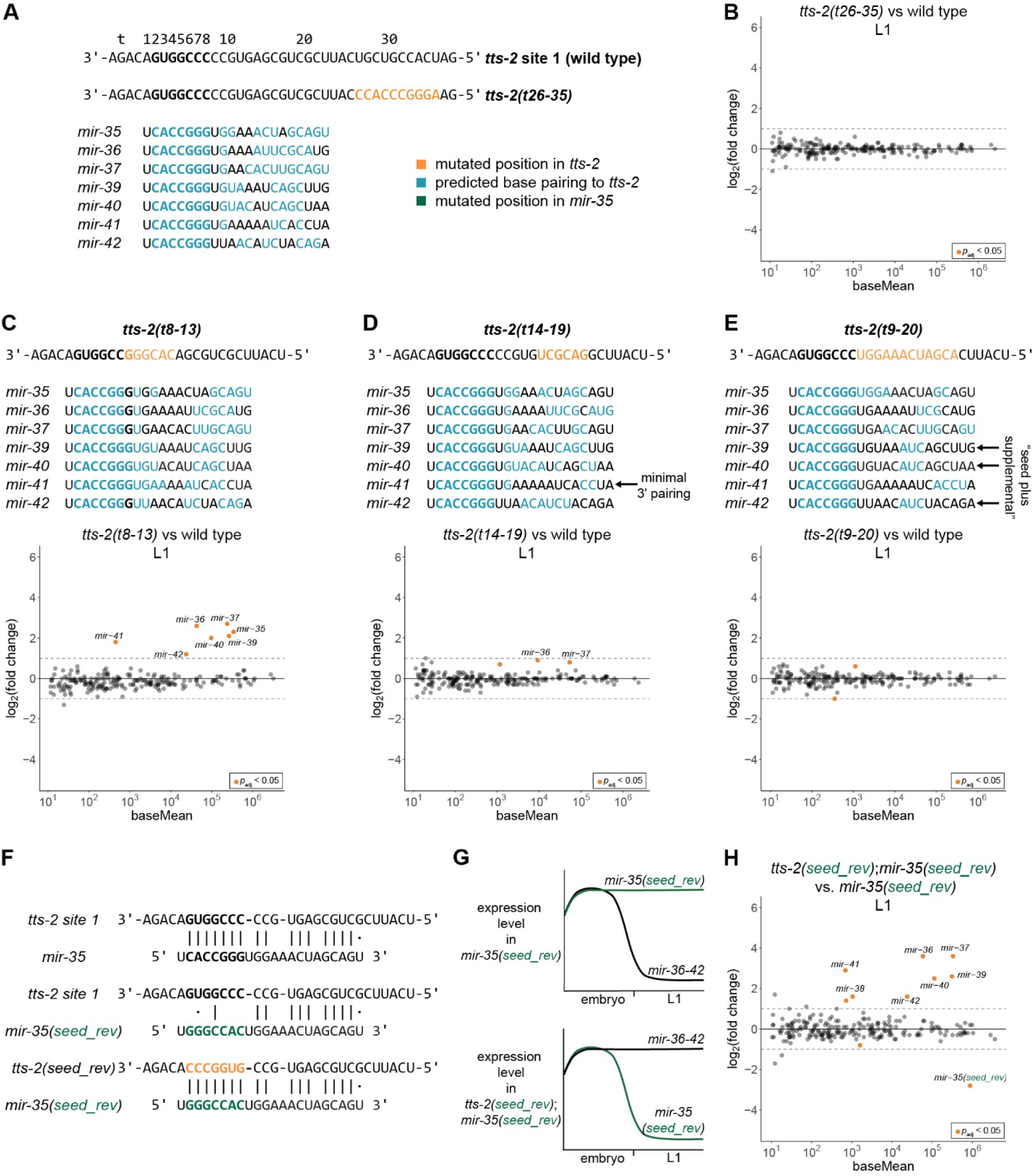
Seed pairing to *tts-2* is sufficient for TDMD. (A) Predicted base pairing between *mir-35-42* family members and wild type *tts-2* or the *tts-2(t26-35)* CRISPR mutant. (B) MA plot of sRNA-seq of *tts-2(t26-35)* vs wild type in L1 animals. (C-E) Additional *tts-2* sequence mutants with predicted *mir-35* family base pairing and sRNA-seq for each mutant in L1 animals. (A, C-E) *mir-38* base pairing is not shown since site 1 mutations do not alter its pairing to redundant site 2. (F) Predicted pairing between wild type and *seed_rev* versions of *mir-35* and *tts-2*. (G) Schematic of the known (top) and expected (bottom) degradation of *mir-35-42* family members in the indicated *tts-2* background. (H) MA plot of sRNA-seq of L1s, comparing *mir-35(seed_rev*);*tts-2(seed_rev)* to *mir-35(seed_rev*) as the control. (B-E, H) sRNA-seq analysis was carried out with three biological replicates per genotype, normalized to total piRNA reads, and miRNAs with baseMean ≥ 10 are shown.

Six additional *tts-2* mutants were assayed by qPCR for their effect on *mir-35* and *mir-36* decay; only one of these [*tts-2(t11-16, 20-25)*] disrupted decay, and qPCR showed that this was due to abrogated expression of the lncRNA (Figure S8B-D). Because overlapping mutations – including t9-20 or t20-25 – do not disrupt decay, it is not immediately clear which portion of the mutation in *tts-2(t11-16, 20-25)* disrupts *tts-2* expression (Figure S8A,D).

Overall, the complement of *tts-2* mutations examined here greatly increases the predicted base pairing conformations between miRNAs and *tts-2* that support efficient decay. These include *mir-41* with *tts-2(t14-19)* which is predicted to have only three non-adjacent paired positions outside the seed (Figure 6D). Another set of examples is *mir-39, -40,* and *-42* with *tts-2(t9-20)*, all of which have only three base pairs in the “supplemental” region often engaged in base pairing in conventional miRNA targets (Figure 6E). These examples, in which EtoL1 decay is intact, demonstrate that *mir-35-42* decay has no specific base pairing requirements outside the seed sequence.

These results together suggest that seed pairing to site 1 of *tts-2* is sufficient to drive miRNA decay. If this is the case, it should be possible to retarget *tts-2* by substituting the seed complement. To test this, we designed a version of *tts-2* in which site 1 binds to a mutant version of *mir-35* that is completely stabilized at the EtoL1 (*mir-35(seed_rev),* Figure 1D, Figure 6F-G)^44^. We targeted this miRNA not only because it escapes EtoL1 decay, but also because its expression from the *mir-35-41* locus ensures that its spatiotemporal expression pattern is compatible with potential *tts-2-* mediated decay. This variant, *tts-2(seed_rev)*, is predicted not only to restore decay of *mir-35(seed_rev)* but also to disrupt decay of endogenous *mir-36-42* since they will no longer bind to the mutated *tts-2* site 1 (Figure 6G, Table S22). We used sRNA-seq to compare *tts-2(seed_rev); mir-35(seed_rev)* to *mir-35(seed_rev)* single mutants in the L1. Indeed, *tts-2(seed_rev)* disrupted decay of wild type *mir-36-42*. Remarkably, *tts-2(seed_rev)* induced potent decay of *mir-35(seed_rev)*, with similar amplitude (7.0-fold) as that induced by the wild type *tts-2/mir-35* pair (7.1-fold change in *tts-2(null)*). We conclude that complementarity to the seed is both necessary and sufficient for *tts-2*-dependent TDMD.

## Discussion

Here we identify a lncRNA that drives TDMD of the essential *mir-35-42* family in a developmentally-timed manner. In contrast to other reported examples of TDMD, *tts-2* driven TDMD relies on pairing between the lncRNA and the miRNA seed region and does not require extensive base pairing to the miRNA 3’ end. Experiments in which the miRNA or the lncRNA is mutated both support this model. Whereas most previously-characterized TDMD triggers distinguish between miRNA family members with differing 3’ ends, *tts-2* drives decay of all members of the *mir-35-42* family without the requirement for pairing beyond the seed. Thus, *tts-2* must use alternative cues, not extensive 3′ end engagement, for EBAX-1 recruitment. This could be a cis element like a specialized RNA structure or also involve trans factors such as an adaptor RNA binding protein. From a biological standpoint, TDMD triggers that drive coherent decay of all members of redundantly-acting miRNA seed families have great regulatory potential for allowing de-repression of the common targets of these families.

The timing of *mir-35-42* decay appears to be patterned by the onset of expression of *tts-2*, whereas EBAX-1 expression starts earlier in embryogenesis^45^. The spatiotemporal patterning of miRNA expression by TDMD triggers is consistent with work in other systems. In mouse tissues, ZSWIM8 is broadly expressed, but tissue-specific subsets of miRNAs undergo ZSWIM8-mediated decay, presumably driven by tissue-specific expression of TDMD trigger RNAs^64,65^. *Transcribed telomerase sequence* may be somewhat of a misnomer for *tts-2* given that it contains only four TTAGGC repeats, and these are in the intron^66^. *tts-2* expression continues beyond embryogenesis according to the GFP knock-in reporter (Figure S5D). Understanding whether *tts-2* has additional functions beyond L1 will be of interest. Intriguingly, *tts-2(null)* animals display overt phenotypes (protruding vulvae) not shared by *tts-2(site_1+2)* or *tts-2(site_1),* suggesting that this abundant transcript may have roles beyond *mir-35-42* TDMD.

To date, identification of TDMD trigger RNAs has proven difficult. Fewer than twenty endogenous TDMD triggers have so far been identified, whereas over a hundred ZSWIM8-sensitive miRNAs (presumed TDMD substrates) have been characterized across human, mouse, zebrafish, *Drosophila*, and *C. elegans* samples^14,19,25–28,31,32,44,46,58,63–65,67^. While the few TDMD triggers known in vertebrates have extensive base pairing to the miRNA 3’ end, relaxed base pairing requirements are observed for some triggers in *Drosophila*^25,28,57^, and here we show that the first reported *C. elegans* TDMD trigger is seed-pairing sufficient. Although bioinformatic approaches have shown some success in identifying trigger RNAs^68^, the minimal requirement for base pairing to *tts-2* suggests that searches for additional TDMD triggers of orphan ZSWIM8-sensitive miRNAs should be broadened to relax base pairing requirements outside the seed.

Interestingly, site 2 of *tts-2* – which acts redundantly with site 1 to induce *mir-38* decay – also expands the repertoire of TDMD trigger sites as the first known site to lack full seed complementarity. While we have not extensively mutagenized it, the impact of site 2 on *mir-38* but not its family members suggests that this site requires extensive 3′ complementarity to compensate for its incomplete seed match. Thus, as more TDMD triggers are characterized, their trajectory roughly characterizes that of conventional targets in revealing a greater diversity of functional base pairing architectures with further study.

The discovery of efficient seed pairing-sufficient TDMD raises the question of what distinguishes miRISC bound to *tts-2* from miRISC bound to conventional targets. The prevailing model of TDMD is that extensive base pairing of the TDMD trigger to the miRNA 3’ end disrupts the binding of the miRNA 3’ end by the Ago PAZ domain, driving a conformational change in Ago that recruits ZSWIM8. Release of the 3′ end is often accompanied by increased prevalence of trimming and tailing (TDTT). Consistent with *mir-35*-*42* TDMD occurring with variable amounts of 3′ end pairing to *tts-2*, we observe TDTT of only *mir-35* and *mir-37*, but not other *mir-35-42* family members. This suggests that display of the miRNA 3ꞌ end outside of the PAZ domain is dispensable for TDMD of most of the family members.

The dispensability of 3’ end display for TDMD of most *mir-35-42* family members suggests that the associated Ago conformational change may also be dispensable. Although most previously-described endogenous triggers of TDMD have a clear requirement for base-pairing to the miRNA 3’ end, whether the basis of this requirement is the resulting conformational change is difficult to test. However, experimental conditions in which Ago PAZ domain mutations force miRNA 3′ end release - and presumably a resulting Ago conformational change – result in miRNA tailing and trimming without driving widespread Ago or miRNA decay^30,69^. An alternative model is that 3’ end pairing is required in many cases of TDMD due to the enhanced affinity it provides, and the Ago conformational change is dispensable. In this case, additional cis features and/or adaptors required to license *tts-2* to drive *mir-35-42* decay may also be required in known instances of the TDMD that require miRNA 3’ end pairing. Some TDMD triggers that require 3’ end pairing also harbor regions of unknown function that are required for TDMD outside the miRNA binding site^22,26,27^, or other topological requirements^70,71^, both of which are consistent with the concept that other unknown features are required to drive ZSWIM8 recruitment.

While TDMD involves ubiquitylation of Ago by the ZSWIM8 E3 ubiquitin ligase, increased Ago levels have not previously been observed upon ZSWIM8 inactivation^31,32^. This is likely because only a small portion of miRNAs expressed in each cell type are substrates of ZSWIM8-mediated TDMD, and therefore, the majority of miRISC pool is not targeted. In contrast, the highly abundant *mir-35-42* family makes up a major portion of all miRNAs expressed in the embryo^41^. Consistent with this and their clearance by TDMD, elevated levels of ALG-1 and ALG-2 are observed in *ebax-1(null)* late embryos compared to wild type. This provides a new layer of support for the model that EBAX-1/ZSWIM8-mediated TDMD does in fact proceed via targeted Ago decay. This observation also shows that EBAX-1 enforces a large-scale developmentally-timed turnover of the overall miRISC pool at the end of embryogenesis. EBAX-1 is unlikely to impact such a large portion of the miRISC pool at later stages of *C. elegans* development wherein TDMD substrates are few and expressed at more moderate levels. Whether ZSWIM8 in other organisms reprograms such a large portion of the miRISC pool in specific developmental stages or tissue compartments remains to be seen. Another remaining open question is the location of the degron in Ago proteins that is recognized and bound by ZSWIM8. Here, N-terminal tags did not disrupt TDMD, suggesting that an N-end degron is not involved. The location of the degron may inform how it is recognized specifically in the context of bound TDMD triggers.

Why *tts-2* is deployed to drive decay of *mir-35-42* is not immediately apparent at the phenotypic level. Animals lacking *tts-2* are superficially wild type, and they do not recapitulate the overt defects of *ebax-1(null)* (egg laying defective, male mating defective), some of which were previously attributed to other EBAX-1 substrates^45^. EBAX-1 and the highly expanded *mir-35* family have recently been implicated in transgenerational induction of predatory mouth-form development in the distantly related *Pristionchus pacificus*^72^. Understanding the role of TDMD of *mir-35-42* in *C. elegans* may require similarly sophisticated experimental paradigms. Nonetheless, the complete conservation of *tts-2* binding site 1 among most species in the *Elegans* supergroup suggests that this site has provided a selective advantage for more than 20 million years.

## Methods

### *C. elegans* husbandry and sample collection

*C. elegans* strains were grown on OP50 as a food source and at 20°C unless otherwise indicated. Bulk embryo samples were collected through standard hypochlorite treatment of synchronized populations 96 hours after feeding L1s^73^. Synchronized L1 samples were collected 24 hours after embryo isolation. Strains used in this study can be found in Table S1, and further allele information in Table S2.

### Total RNA isolation

RNA was extracted using TRIzol Reagent (Life Technologies). RNA extraction was carried as previously described with minor modification^74^. After TRIzol addition, samples underwent three freeze-thaw cycles on dry ice and were vortexed at room temperature for 15 minutes prior to addition of chloroform. The resulting aqueous phase was isolated and extracted once with an equal volume of 25:24:1 phenol:chloroform:isoamyl alcohol, pH=4.5. This aqueous phase was transferred to a fresh tube and an equal volume of isopropanol was added alongside 1μL of GlycoBlue (Thermo Fisher Scientific). Samples were stored at -80°C until frozen to promote RNA precipitation, then thawed on ice. Samples were washed twice with 1ml of 80% ethanol, and 20μL of RNase-free water was used for re-suspension.

### CRISPR/Cas9-mediated genome editing

Guide RNAs were chosen for gene editing based upon previous used design criteria, favoring both high GC content and CRISPRScan scores where possible^75^. Homologous repair templates were designed with 35-bp homology arms to promote homology directed repair. Alt-R crRNAs and tracrRNA (IDT) were annealed in IDT Duplex Buffer at a final concentration of 10µM by heating to 95C for five minutes, then cooling at room temperature for 5 minutes, before placing on ice. Annealed guide RNAs were 4µM (total of all guides) in injection mixes. For gene editing co-injections markers, see details in Table S2.

### Chimeric eCLIP Library Preparation for C. elegans Samples

Chimeric eCLIP experiments were adapted for *C. elegans* based on the protocol described by Manakov *et al.*^55^ See supplemental methods all modifications and details. Animals were grown in standard liquid culture for 96h from L1, followed by harvest of embryos by hypochlorite treatment^73^. Embryos were spread on unseeded NGM plates and UV cross-linked at 254nm via a Stratalinker set at energy setting 3000 (3kJ/m^2^). Samples were snap frozen on dry ice and stored at -80°C. Pellets were resuspended in 1mL of homogenization buffer [100mM NaCl, 25mM HEPES pH 7.5,, 250µM EDTA, 0.10% NP-40, 2mM DTT, 25U/ml RNAse inhibitor (Ribolock), 1x cOmplete mini protease inhibitor (Roche)]. Sample were lysed using a Bioruptor, with 30 cycles of 30s on, 30s off. Protein was quantified by BCA, normalized to 3mg/mL, then stored at -80°C until use. Samples containing 3 mg total protein were incubated with 50 μL of anti-FLAG magnetic beads (Sigma, M8823) with gentle rotation at 4°C overnight.

To ligate chimeric RNA duplexes, T4 RNA Ligase 1 (NEB, M0437) was added, and reactions were incubated overnight at 4 °C. After additional wash steps, 3′ RNA adaptors (/5rApp/NNNNTGGAATTCTCGGGTGCCAAGG/3ddC/) were conjugated overnight at 16 °C using T4 RNA Ligase 2 truncated K227Q mutant (NEB, M0351). The RNA–protein complexes were resolved by SDS–polyacrylamide gel electrophoresis, and regions corresponding to 100–250 kDa, representing the ALG-2–miRNA–target complexes, were excised from the membrane and treated with proteinase K to release ligated RNA molecules. Reverse transcription was performed using SuperScript IV (Invitrogen, 18090050) with RT primers (GCCTTGGCACCCGAGAATTCCA). Amplification of cDNA libraries was carried out with 12 PCR cycles, and products ranging from 180 bp to 400 bp were enriched by size selection on 8% native polyacrylamide gels. See supplemental methods for details. Libraries were sequenced on Illumina NovaSeq platforms.

### Bioinformatic Processing and Analysis of Chimeric eCLIP Data

Sequencing reads were processed through a multi-step computational workflow. The *C. elegans* transcriptome reference was obtained from Ensembl (WBcel235) to construct a comprehensive transcript database used by Hyb^76^. Raw sequencing data were subjected to adapter trimming using Cutadapt v2.10^78^ with the parameters -a TGGAATTCTCGGGTGCCAAG -A GATCGTCGGACTGTAGAACT. Paired-end reads were merged using PEAR v0.9.6^80^, followed by deduplication with fastx_collapser v0.0.14 to remove PCR duplicates. Unique molecular identifiers (UMIs) were trimmed using Cutadapt (-u 4 -u -4 -m 18) to retain high-confidence fragments.

Processed reads were analyzed with Hyb to identify miRNA–target chimeric interactions, which generated intermediate output files in the .viennad and .hyb formats. Custom Python scripts (CLASH.py) were used to parse and integrate information from these files, extract all miRNA– target pairs, and normalize miRNA–target counts to the total number of hybrids. The CLASH.py script is available at https://github.com/UF-Xie-Lab/TDMD-in-Celegans.

### Single molecule fluorescence in situ hybridization

For single molecule fluorescence in situ hybridization (smFISH), embryos were scraped from the plate and subjected to freeze-crack on 0.01% poly-lysine coated slides, followed by fixation in - 20°C methanol overnight. *In situ* protocol was followed as in Scholl et al., 2024^77^.

Microscopy was performed using a Zeiss Axio Observer equipped with a CSU-W1 SoRA spinning disk scan head (Yokogawa) and an iXon Life 888 EMCCD camera (Andor) using a 63x objective with a 2.8x relay lens (Yokogawa). Images were taken using Slidebook software (Intelligent Imaging Innovations).

### Airlocalize analysis

High resolution smFISH images were analyzed using Airlocalize^79^. Entire Z-stack *in situ* images were processed in Fiji by applying a background subtraction of 50 pixels. All images from a single experiment were manually thresholded together. To identify intensities corresponding to single mRNA molecules, images of bean stage embryos were used. PSF Width and Detection Threshold were initially set interactively, and 5 foci were selected and fitted with a gaussian curve to set the PSF. Threshold was adjusted until all foci were detected by the program. The same PSF and threshold value was used on all images taken on the same day using batch mode. The median of the Integrated Intensity (output from Airlocalize) for all bean stage images was assigned as the intensity of a single mRNA molecule. That intensity value was used to bin foci corresponding to 2, 3, or more molecules. Python codes used for processing Airlocalize data are deposited on Github: https://github.com/yliu380/AirLocalize_Data_Process

### Small RNA sequencing library preparation and data analysis

Small RNA libraries were prepared through a modified protocol using the NEBNext Small RNA kit (NEB). Synthetic RNA spike-ins were added to 600ng of input RNA per sample prior to library preparation. Following cDNA synthesis, size selection of products between 65-75nt was carried out using an 8% urea gel (National Diagnostics). Final library PCR preparation was carried out with 15 cycles followed by purification using the NEB Monarch PCR purification kit. Data analysis was carried out using the NIH High Performance Computing cluster. Adaptors were trimmed using cutadapt (ver. 5.0) with the -q 20 flag. Reads were mapped to the *C. elegans* genome build WS280 modified to include an artificial chromosome for mutant microRNAs and synthetic spike-ins^44^. Mapping was performed using bowtie2 (ver. 2.5.3) with --no-unal --end-to-end – sensitive -U flags. BAM files were then sorted and indexed using samtools (ver 1.21). Read counts were generated for miRNAs and piRNAs using htseq (ver. 2.0.4) with --mode union --nonunique fraction -a 0 flags. Counts for miRNAs were assigned using a modified gff file from mirGeneDB^81^. piRNA counts were assigned using a gff file obtained from https://www.pirnadb.org/^82^. Differential analysis was carried out using DEseq2 with size factors specified using total mapped reads, piRNA or spike-ins for normalizations as indicated in corresponding figure legends^83^. pAdj was calculated using the DEseq2 p-value corrected for multiple comparisons using the “hochberg” correction. Tailing and trimming analysis was performed using miTRATA ^84^. Detailed information about the samples used for small RNA-seq can be found in Table S3 and raw counts in Table S4.

### Taqman miRNA quantitative Polymerase Chain Reaction (qPCR)

Quantification of miRNAs was carried out using miRNA Taqman qPCR. In brief, 10ng of input total RNA (concentration 6ng/μL) was used to perform 5μL reverse transcription reactions using the TaqMan MicroRNA Reverse Transcription kit (ThermoFisher) with miRNA-specific probes. RT products were diluted 1:4 in dH_2_0. Technical triplicate qPCRs contained 1.33μL of diluted RT product in 5μL reactions using Taqman Universal Mastermix II with UNG kit (ThermoFisher) and were run on an Applied Biosystems QuantStudio 6 instrument. The 2^−Δ Δ Ct^ method was used for analysis. Absolute quantification of *mir-35* was carried out using synthetic *mir-35* (Dharmacon).

### Real-time qPCR

The KAPA SYBR FAST One-Step qRT-PCR Master Mix (2X) Kit (Fisher Scientific) was for qPCR according to the manufacturer’s instructions on an Applied Biosystems QuantStudio Pro 6. An input of 10ng RNA was used per 20μL reaction mix, and the reaction was split into triplicate 5μL reactions before running on the instrument. Data was analyzed using the 2^−Δ Δ Ct^ method. Absolute quantification of *tts-2* was carried out using in vitro transcribed *tts-2* generated using the HiScribe T7 Quick High Yield RNA Synthesis Kit (NEB) according to the kit’s protocol. Oligonucleotides used for qPCR can be found in Table S2.

### Fluorescent protein image quantification

Cell profiler was used to semi-automate embryo image quantification from wide field images acquired using an Andor Zyla cMos camera and 20x water immersion objective on a Nikon eclipse Ni microscope. See supplemental methods for details of Cell Profiler workflow. Representative images shown were acquired using a Nikon C2 confocal.

### RNA-RNA pairing prediction

Interactions between the *mir-35-42* and *tts-2* variants were predicted using the ViennaRNA package (ver. 2.5.1) ^85,86^. RNAduplex was utilized with a temperature set using the flag -T ‘20’, to represent typical *C. elegans* growth conditions. The input *mir-35-42* sequences began at the first position of the seed sequence (nt 2) and input *tts-2* sequences were the predicted interaction site with additional 10nt downsteam and upstream.

### Statistical analysis

Statistical analysis was carried out in Graphpad Prism 10 software where applicable.

## Data accessibility

All deep sequencing datasets have been deposited at the NCBI GEO under the accession number GSE303817. Reviewers may access the data until release using token “yvipssoknhktnmp”.

## Acknowledgements

Many strains used in this research were received from Yishi Jin and the CGC, which is funded by the NIH Office of Research Infrastructure Programs (P40 OD010440). This work used the computational resources of the NIH HPC Biowulf cluster (https://hpc.nih.gov). The NCI CCR Genomics core performed most of the deep sequencing. WormBase was extensively used in the design and execution of this study^59^.

Thank you to Shawn Ahmed and members of the McJunkin lab and Baltimore Worm Club for helpful discussions and to Helge Grosshans and Rajani Gudipatti for sharing unpublished data. This work was funded by the NIDDK Intramural Research Program (ZIADK075147).

This research was supported by the Intramural Research Program of the National Institute of Diabetes and Digestive and Kidney Diseases (NIDDK) within the National Institutes of Health (NIH). The contributions of the NIH author(s) were made as part of their official duties as NIH federal employees, are in compliance with agency policy requirements, and are considered Works of the United States Government. However, the findings and conclusions presented in this paper are those of the author(s) and do not necessarily reflect the views of the NIH or the U.S. Department of Health and Human Services.

