## Supplemental Figures for "A lncRNA drives developmentally-timed decay of all members of an essential microRNA family"

**List of Supplemental Materials**

Figure S1. ALG-2 loading is neither sufficient nor necessary for *mir-35-42* decay at EtoL1.
Figure S2. Summary of chimeric eCLIP hybrid reads.
Figure S3. SL1 trans-splicing of *tts-2*.
Figure S4. *tts-2* smFISH signal is specific.
Figure S5. GFP knock-in at the *tts-2* locus shows late embryonic expression and continued transcription in larvae and adults.
Figure S6. Trimming and tailing of *mir-35-42* family members.
Figure S7. Absolute quantification of *mir-35* and *tts-2*.
Figure S8. Quantification of *tts-2* transcript and *mir-35* and *mir-36* in additional *tts-2* mutant backgrounds by qPCR.
Supplemental Methods
Table S1. Strains used in this study.
Table S2. Alleles generated and oligonucleotides used in this study.
Table S3. Samples used for deep sequencing.
Table S4. Small RNA deep sequencing raw feature counts of all genome-mapped reads.
Table S5. *ebax-1(null)* vs. wild type DESeq2 output, embryo, whole library normalized.
Table S6. *ebax-1(null)* vs. wild type DESeq2 output, L1, whole library normalized.
Table S7. *alg-1(null)* vs. wild type DESeq2 output, embryo, spike-in normalized.
Table S8. *alg-1(null)* vs. wild type DESeq2 output, L1, spike-in normalized.
Table S9. *alg-2(null)* vs. wild type DESeq2 output, embryo, spike-in normalized.
Table S10. *alg-2(null)* vs. wild type DESeq2 output, L1, spike-in normalized.
Table S11. Normalized CLASH hybrids.
Table S12. *tts-2(null)* vs. wild type DESeq2 output, L1, piRNA normalized.
Table S13. *ebax-1(null)* vs. wild type DESeq2 output, L1, piRNA normalized.
Table S14. *ebax-1(null); tts-2(null)* vs. wild type DESeq2 output, L1, piRNA normalized.
Table S15. *tts-2(site1+2_del)* vs. wild type DESeq2 output, L1, piRNA normalized.
Table S16. *tts-2(site1_del)* vs. wild type DESeq2 output, L1, piRNA normalized.
Table S17. *tts-2(site2_del)* vs. wild type DESeq2 output, L1, piRNA normalized.
Table S18. *tts-2(t26-35)* vs. wild type DESeq2 output, L1, piRNA normalized.
Table S19. *tts-2(t8-13)* vs. wild type DESeq2 output, L1, piRNA normalized.
Table S20. *tts-2(t14-19)* vs. wild type DESeq2 output, L1, piRNA normalized.
Table S21. *tts-2(t9-20)* vs. wild type DESeq2 output, L1, piRNA normalized.
Table S22. *mir-35(seed_rev); tts-2(seed_rev)* vs. *mir-35(seed_rev)* DESeq2 output, L1, piRNA normalized.

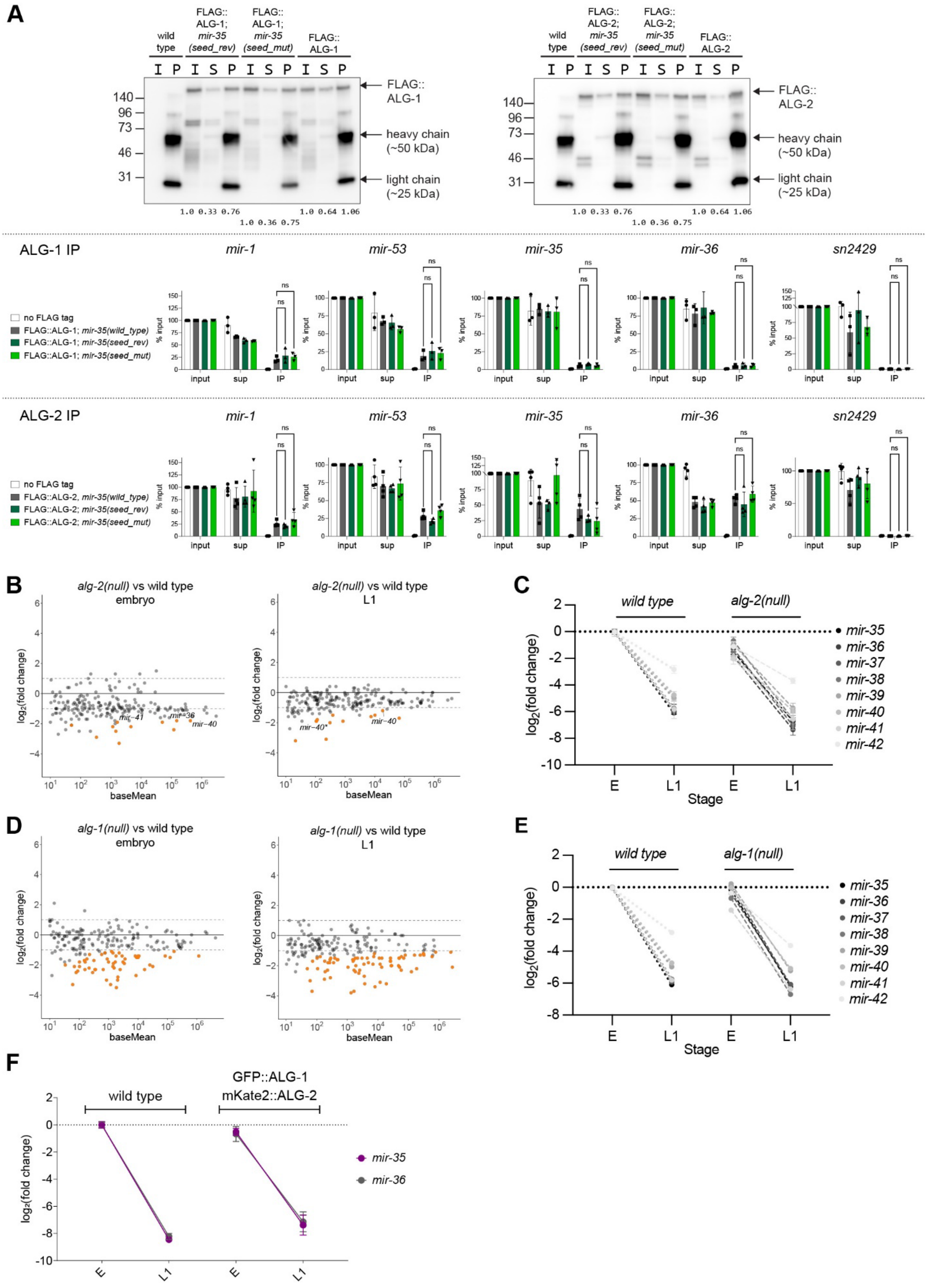
A
wild type
FLAG:: ALG-1; mir-35 (seed_rev)
FLAG:: ALG-1; mir-35 (seed_mut)
FLAG:: ALG-1
FLAG:: ALG-1
heavy chain (~50 kDa)
light chain (~25 kDa)
wild type
FLAG:: ALG-2; mir-35 (seed_rev)
FLAG:: ALG-2; mir-35 (seed_mut)
FLAG:: ALG-2
FLAG:: ALG-2
ALG-1 IP
mir-1
mir-53
mir-35
mir-36
sn2429
no FLAG tag
FLAG::ALG-1; mir-35(wild_type)
FLAG::ALG-1; mir-35(seed_rev)
FLAG::ALG-1; mir-35(seed_mut)
ALG-2 IP
FLAG::ALG-2, mir-35(wild_type)
FLAG::ALG-2; mir-35(seed_rev)
FLAG::ALG-2; mir-35(seed_mut)
B
alg-2(null) vs wild type embryo
alg-2(null) vs wild type L1
baseMean
log2(fold change)
C
wild type
alg-2(null)
Stage
D
alg-1(null) vs wild type embryo
alg-1(null) vs wild type L1
E
wild type
alg-1(null)
F
wild type
GFP::ALG-1 mKate2::ALG-2
mir-35
mir-36

**Figure S1. ALG-2 loading is neither sufficient nor necessary for *mir-35-42* decay EtoL1.** (A) Top: Representative western blots of FLAG::ALG-1 (left) and FLAG::ALG-2 (right) immunoprecipitations. I, Input; S, Supernatant; P, Immunoprecipitation. Five percent of each was loaded. Numbers below blot are proportion of input FLAG::ALG signal recovered in supernatant or immunoprecipitation. Bottom: miRNA Taqman qPCR was performed to assess the relative percent recovery in each fraction of the IP. miRNAs loaded into ALG-1 and ALG-2 (*mir-1* and *mir-53*), miRNAs preferentially loaded into ALG-2 (*mir-35* and *mir-36*), and non-miRNA control (sn2429) are shown. The *mir-35* seed mutants are preferentially loaded into ALG-2, similarly to wild type. (B, D) MA plots of sRNA-seq in indicated genotypes and stages. Three biological replicates, normalized to spike-ins. miRNAs with baseMean>10 are shown. (C, E) miRNA levels normalized to wild type embryo levels (sRNA-seq). (F) miRNA Taqman qPCR in wild type or fluorescently-tagged Ago strain. Three biological replicates, normalized to sn2429, then to wild type embryo levels.

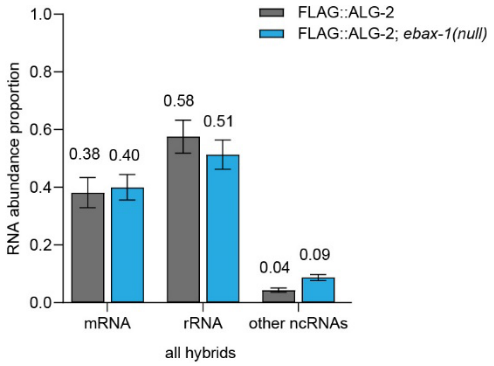


**Figure S2. Summary of chimeric eCLIP hybrid reads.** Comparison of RNA classes in miRNA-containing hybrids from embryonic ALG-2 chimeric eCLIP in wild type or *ebax-1(null)*.

| Total RNAseq reads near *tts-2* 5' end (trimmed to show end sequence) | number of reads | |
|---|---|---|
| G | 2 | ambiguous reads near 5' end |
| TG | 1 | |
| GTG | 47 | |
| TGTG | 27 | |
| GATGTG | 2 | |
| GAGATGTG | 16 | 98.9% SL1 trans-spliced reads |
| TGAGATGTG | 38 | |
| TTGAGATGTG | 40 | |
| TTTGAGATGTG | 14 | |
| GTTTGAGATGTG | 261 | |
| AGTTTGAGATGTG | 2 | |
| AAGTTTGAGATGTG | 3 | |
| CAAGTTTGAGATGTG | 34 | |
| CCAAGTTTGAGATGTG | 32 | |
| CCCAAGTTTGAGATGTG | 3 | |
| TTTCAGATGTG | 1 | 1.1% unspliced reads |
| TATGATTTCAGATGTG | 1 | |
| TTTTTTTGATTACTGTAGTATTATGATTTCAGATGTG | 1 | |
| TTTTTTTGATTACTGTAGTCTTATGATTTCAGATGTG | 1 | |
| AAAATATTTTTTTTGATTACTGTAGTCTTATGATTTCAGATGTG | 1 | |

**Figure S3. SL1 trans-splicing of *tts-2*.** Reads spanning the 5′ end of *tts-2* show efficient addition of SL1 splice leader. Reads were obtained from total RNA seq from wild type embryo samples (Kotagama, et al. 2024).

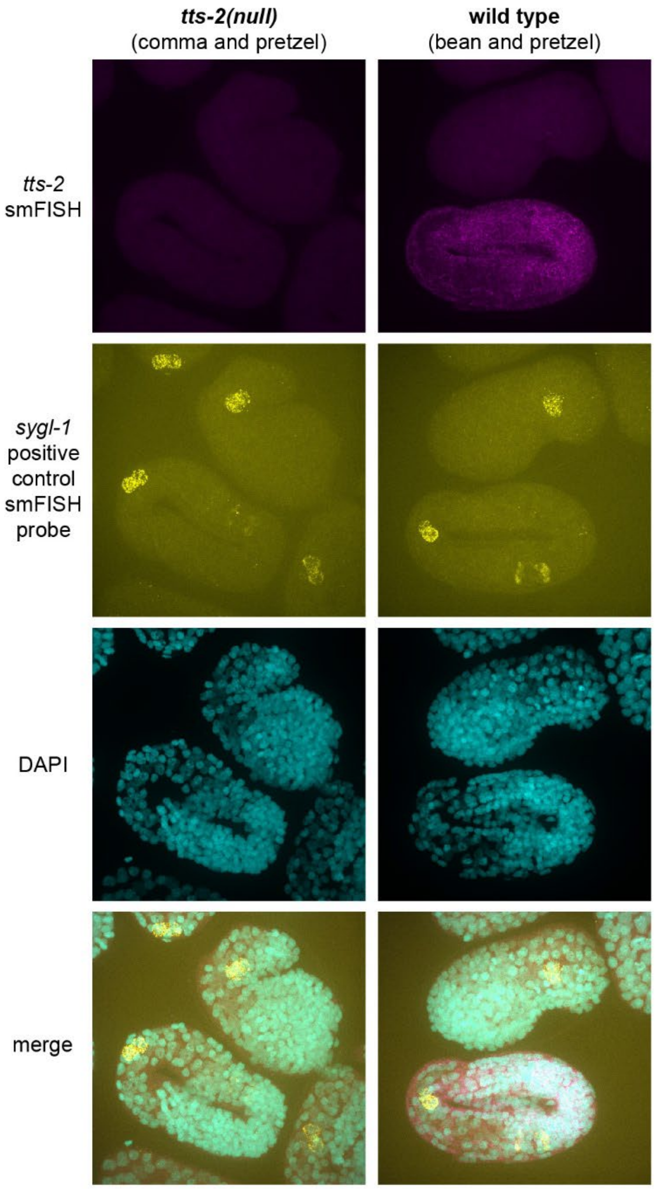


**Figure S4.** ***tts-2*** **smFISH signal is specific.** Representative images of *tts-2* smFISH signal in *tts-2(null)* (left) or wild type (right). An unrelated smFISH probe (*sygl-1*) serves as a positive control signal present in both genotypes.

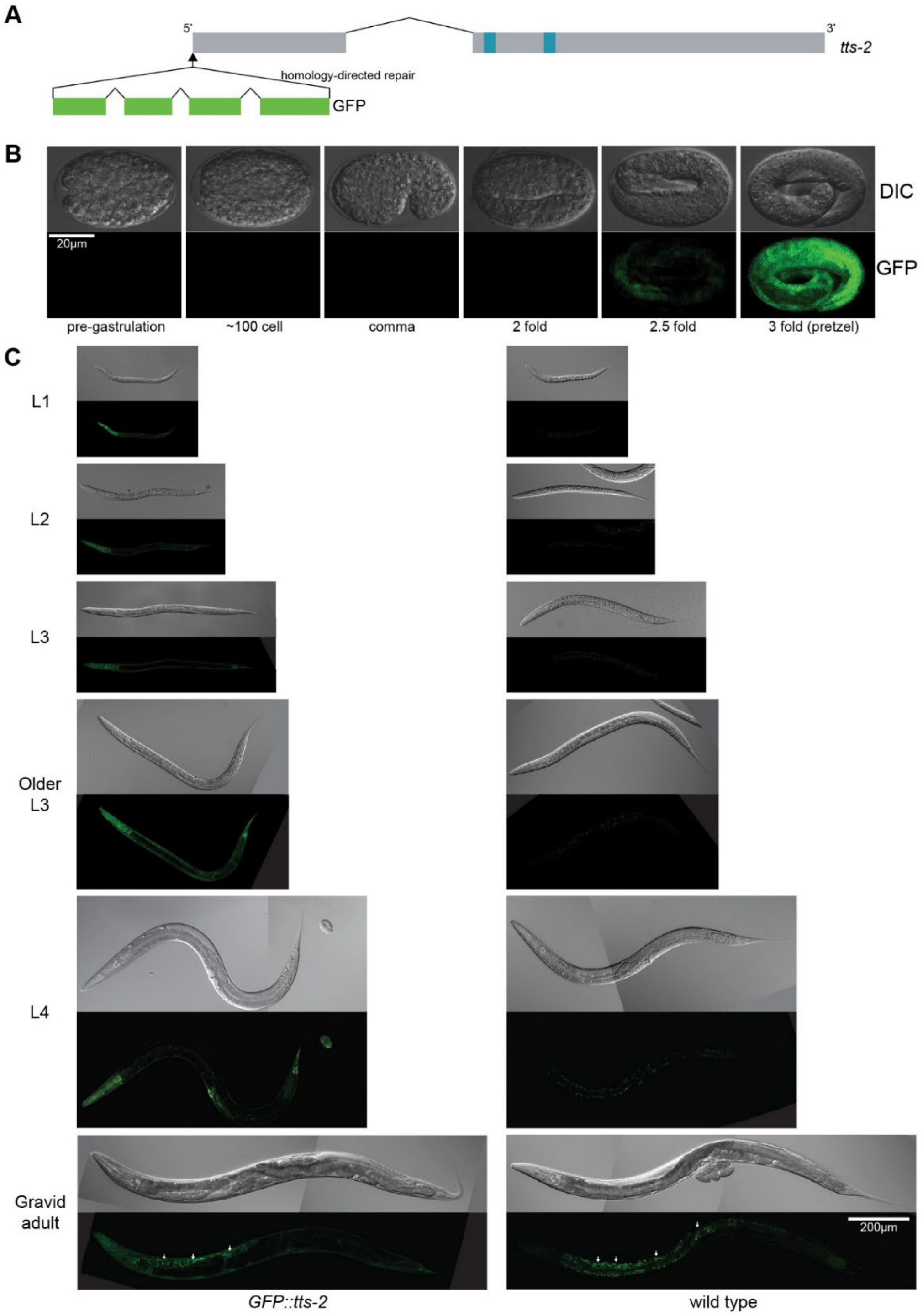


**Figure S5. GFP knock-in at the *tts-2* locus shows late embryonic expression and continued transcription in larvae and adults.** (A) Schematic of GFP knock-in at the 5′ end of the *tts-2* locus. The *tts-2* sequence is fused to the GFP CDS as its 3′ UTR. (B) Expression of GFP::*tts-2* is detected in late embryos. (C) Expression continues in larvae and adults with notable expression in the head throughout development, bright vulva expression in L4, and visible body wall expression in L3 through adult. Stage-matched wild type animals are shown to illustrate levels of autofluorescence. Intestinal autofluorescence is marked by arrows in adult images.

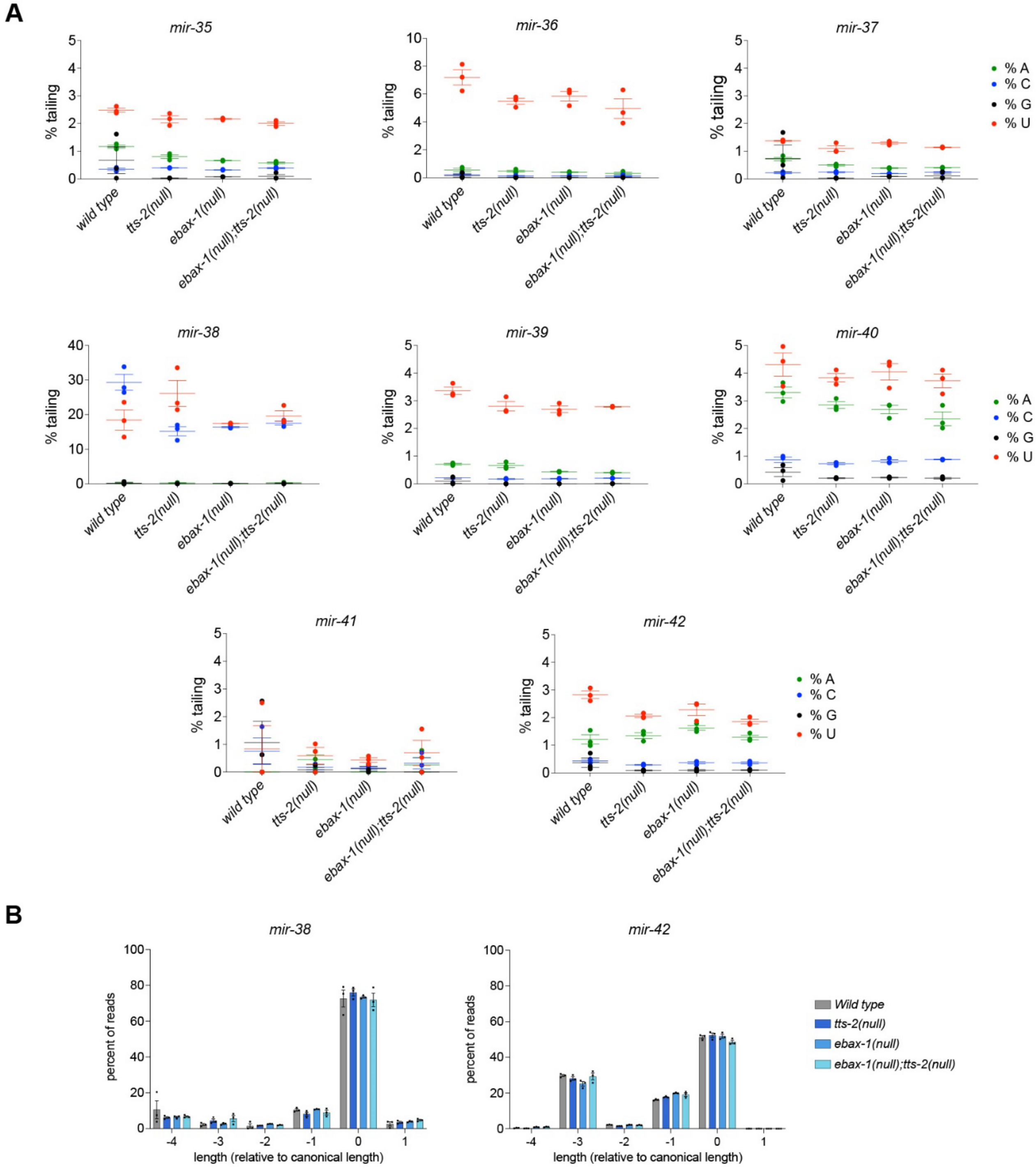


**Figure S6. Trimming and tailing of *mir-35-42* family members.** (A) Percent tailed isoforms in L1s of indicated genotypes. (B) Trimming of *mir-38* and *mir-42* in L1s of indicated genotypes. (A-B) Three biological replicates are shown with mean and SEM.

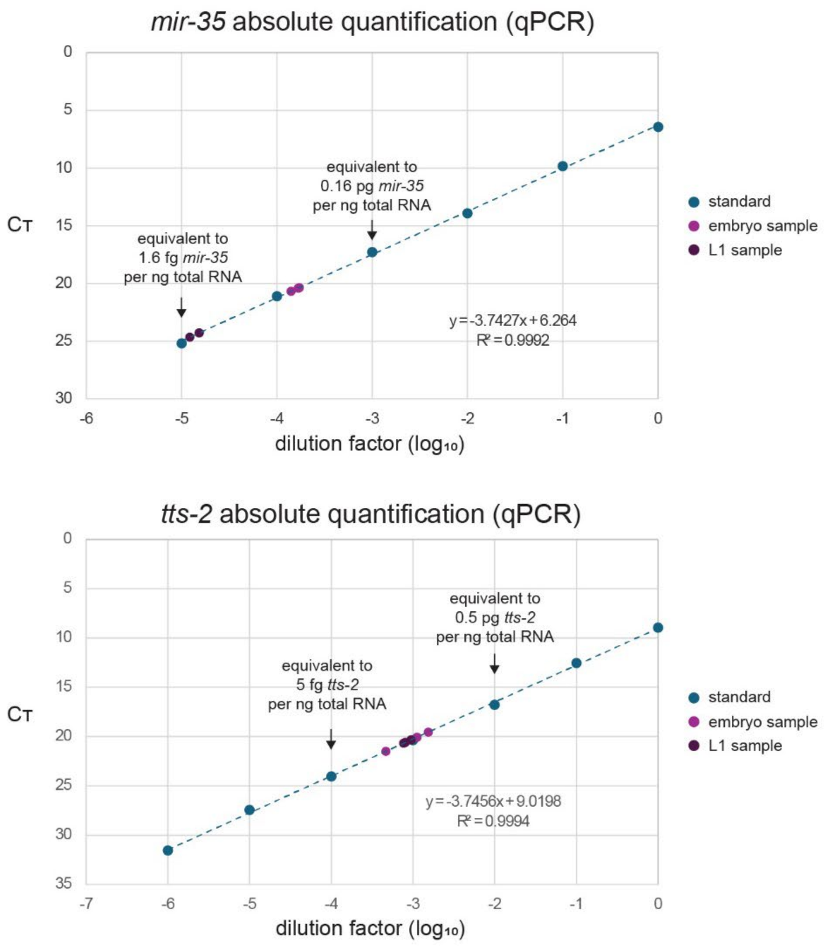


**Figure S7. Absolute quantification of *mir-35* and *tts-2*.** Plots show standard curves for *mir-35* and *tts-2* generated using an RNA oligonucleotide for *mir-35* or in vitro transcribed *tts-2*. Three biological replicate embryo and L1 samples were quantified, and their interpolated positions on the standard curve are shown.

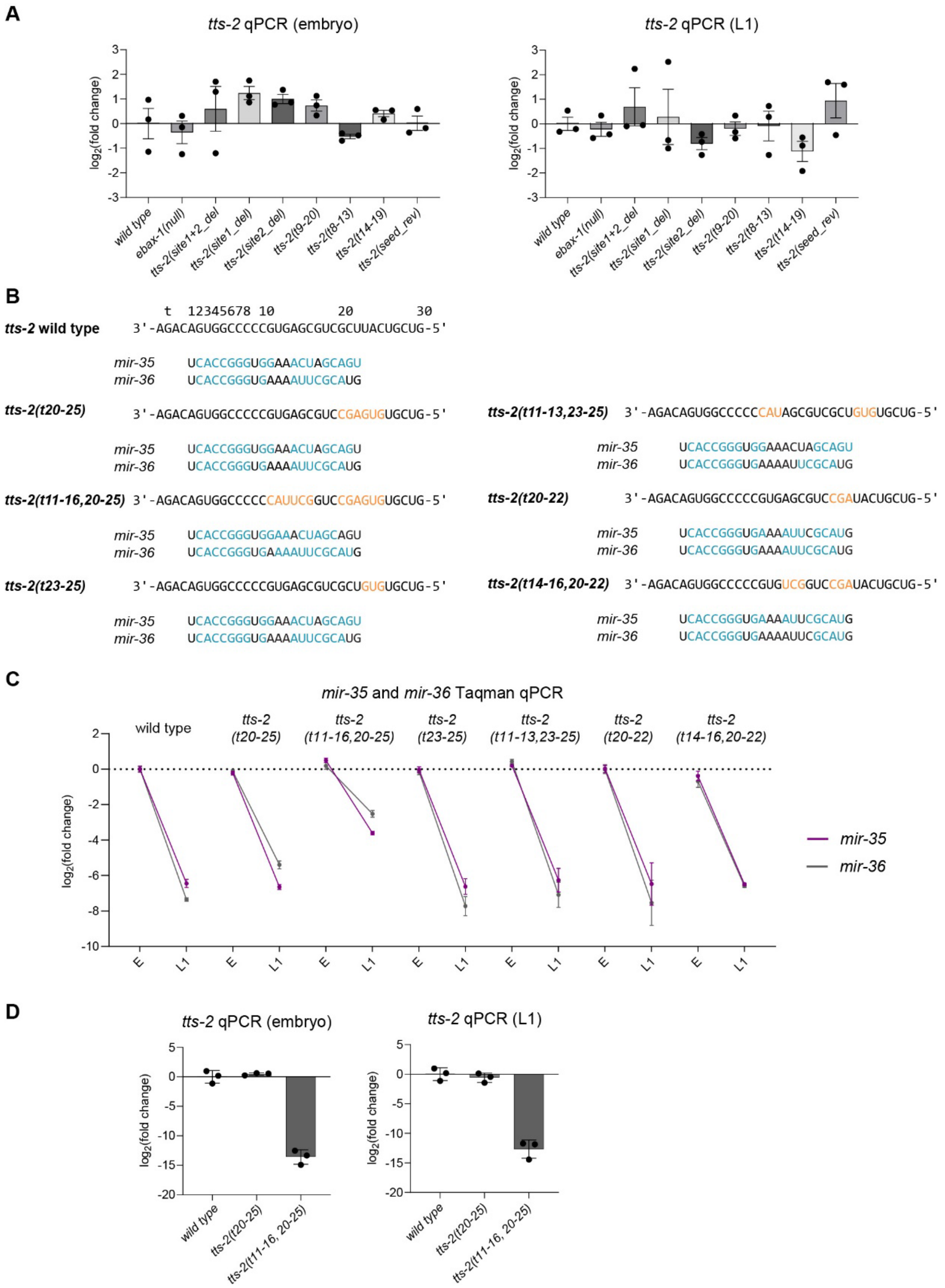

**Figure S8. Quantification of *tts-2* transcript and *mir-35* and *mir-36* in additional *tts-2* mutant backgrounds by qPCR.** (A, D) RT qPCR quantification of *tts-2* in wild type or backgrounds with mutations in *ebax-1* or *tts-2*. Mean and SEM of three biological replicates shown. (B) Predicted base pairing of *mir-35* and *mir-36* with *tts-2* site 1 mutations shown. Only *mir-35* and *mir-36* are shown since other miRNAs were not quantified. (C) miRNA Taqman qPCR of *mir-35* and *mir-36* in indicated *tts-2* mutant backgrounds. Mean and SEM of three biological replicates shown.

**Supplemental Methods**

**Semi-automated fluorescence quantification by Cell Profiler**

Images were processed in batches of no more than 40 images to prevent software crashes.

The following steps were used to process DIC images to sharpen edges for embryo recognition: EnhanceOrSuppressFeatures/Suppress/Feature size 3 > EnhanceOrSuppressFeatures/Enhance/ Smoothing Scale 2.0, Shear angle 225, Decay 0.85. These settings must be optimized for each imaging session to maximize the number of embryos correctly identified in the DIC channel.

Next, embryos were identified: IdentifyPrimaryObjects/Use advanced settings/Typical diameter of objects Min 75 Max 175/Discard objects outside diameter range/Discard objects touching the border of the image/Threshold strategy Global/Thresholding method Robust Background/Lower outlier fraction 0.05/Upper outlier fraction 0.05/Averaging Method Mean/Variance Method Standard deviation/# of deviations 2.2/Threshold smoothing scale 0/Threshold correction factor 1/Lower and upper bounds on threshold 0.0 1/Method to distinguish clumped objects None/Fill holes in identified objects After both thresholding and declumping/Handling of objects if excessive number of objects identified Continue.

A mask of the identified objects was created: FillObjects/Name of output smoothed_objects/Filling method Convex hull > MaskImage/Select the input image DIC/Use objects or image as a mask Objects/Invert the mask? No. > SaveImages/tiff/Save with lossless compression Yes/Every cycle.

The resulting images with object masks were used to manually curate the automated embryo identification. Non-embryo debris, truncated embryos, and clumped embryos were deleted from dataset (at the level of downstream intensity measurements). Embryo stage for each retained object was also manually scored using these images.

Intensity measurements were made: MeasureObjectIntensity/GFP/mCherry/smoothed_objects > ExportToSpreadsheet.

**Chimeric eCLIP Library Preparation for C. elegans Samples**

| | |
|---|---|
| iCLIP Lysis Buffer | 50 mM Tris-HCl (pH 7.4), 100 mM NaCl, 1% (v/v) NP-40, 0.1% (w/v) SDS, and 0.5% (w/v) sodium deoxycholate (light-sensitive) and 1X protease inhibitor cocktail (added fresh) |
| High Salt Wash Buffer | 50mM Tris-HCl (pH 7.5), 1M NaCl, 1mM EDTA, 1% NP-40, 0.1% SDS, and 0.5% sodium deoxycholate (light-sensitive). |
| Wash Buffer | 20mM Tris-HCl (pH 7.5), 10 mM MgCl2, 0.2% Tween-20 and 5mM NaCl. |
| 1X PNK7 Buffer | 70 mM Tris-HCl (pH 7), 10 mM MgCl2 |
| 10X PNK7 Buffer | 700 mM Tris-HCl (pH 7), 100 mM MgCl2 |
| T4 PNK Minus Mix | 84.5 µL $H_2O$, 10 µL 10X PNK pH 7 buffer, 1 µL 0.1 M ATP, 0.5 µL 4 M NaCl, 1 µL Murine RNase Inhibitor, 3 µL T4 PNK Minus enzyme (NEB, M0236) |
| Intermolecular Ligation Mix | 80.4 µL $H_2O$, 18 µL 10X T4 RNA Ligase buffer, 3.6 µL 1% Tween-20, 5.4 µL DMSO, 1.8 µL ATP (100 mM), 54 µL PEG8000 (50%), 2.4 µL Murine RNase Inhibitor, 14.4 µL T4 RNA Ligase I (NEB, M0437, high concentration) |
| FastAP mix | 40 µL $H_2O$, 5 µL 10X FastAP buffer, 2 µL Murine RNase Inhibitor, 3 µL FastAP enzyme |

| | |
|---|---|
| PNK mix | 126 µL $H_2O$, 20 µL 10X PNK buffer, 4 µL T4 PNK enzyme |
| miRCat-33 | /5rApp/NN NNT GGA ATT CTC GGG TGC CAA GG/3ddC/ |
| 3' adapter ligation mix | 42 µL $H_2O$, 8 µL 10X RNA ligase buffer, 16 µL PEG8000 (50%), 2 µL Murine RNase Inhibitor, 8 µL miRCat-33 3′ linker (10 µM), 4 µL T4 RNA Ligase 2 truncated K227Q |
| Proteinase K SDS mix | 130 µL PKS Buffer, 20 µL Proteinase K (20 mg/mL) |
| RTP primer | GCCTTGGCACCCGAGAATTCCA |
| 4X Mn SSIII/IV buffer | 200 mM Tris-HCl (pH 8), 300 mM KCl, and 12 mM $MnCl_2$ |
| RT mix | 5 µL 4X Mn SSIII/IV buffer, 2.8 µL $H_2O$, 1 µL 0.1 M DTT, 0.4 µL Murine RNase Inhibitor, 0.8 µL SuperScript IV (Invitrogen, 18090050) |
| RLTW Buffer | 1x Qiagen, #79216; 0.025% Tween-20 |
| TT Elution Buffer | 10 mM Tris-HCl (pH 7.5), 0.01% Tween-20, and 0.1 mM EDTA. |
| RA5 DNA adapter | /5Phos/NNNNGATCGTCGGACTGTAGAACTCTGAAC/3SpC3/ |
| 5' adapter ligation mix | 1.15 µL $H_2O$, 1 µL 10X RNA ligase buffer, 0.2 µL 0.1 M DTT, 0.1 µL 0.1 M ATP, 0.2 µL Tween-20, 3.6 µL PEG8000 (50%), 1 µL RNA Ligase high concentration (NEB, M0437), and 0.3 µL 5′ Deadenylase (NEB, M0331) |

For chimeric eCLIP, each sample was thawed and treated with 10 µL of Turbo DNase (Invitrogen, AM2239) and 20 µL of RNase I diluted 1:100, followed by incubation at 37°C for 5 minutes with shaking at 1200 rpm. After digestion, lysates were clarified by centrifugation at 21,000 × g for 10 minutes at 4°C.

Immunoprecipitation was carried out by incubating the supernatant with 50 µL of anti-FLAG magnetic beads (Sigma, M8823) specific for ALG2-FLAG-expressing C. elegans samples. The incubation proceeded overnight at 4°C with constant rotation. Beads were then washed three times with 500 µL of ice-cold High Salt Wash Buffer, followed by three washes with 500 µL of Wash Buffer, and once with 200 µL of cold 1× PNK buffer.

For phosphorylation on beads, 100 µL of T4 PNK Minus Mix was added and incubated at 37°C for 20 minutes. This was followed by a single wash with 500 µL High Salt Wash Buffer and three additional washes with 500 µL Wash Buffer. Intermolecular ligation was performed by incubating the beads overnight at 4°C in 180 µL of Intermolecular Ligation Mix. Post-ligation, beads were washed three times each with 500 µL of cold High Salt Wash Buffer and Wash Buffer.

To dephosphorylate the RNA, beads were incubated with 50 µL of FastAP mix at 37°C for 10 minutes. Without removing the FastAP mix, 150 µL of PNK mix was added directly and incubated again at 37°C for 20 minutes. Following this step, the beads were washed once with 500 µL of High Salt Wash Buffer and three times with 500 µL of Wash Buffer.

For 3′ linker ligation, the beads were resuspended in 80 µL of 3′ adapter ligation mix and incubated overnight at 16°C with intermittent shaking (15 seconds every minute at 1400 rpm). After ligation, the beads were sequentially washed with 500 µL of cold Wash Buffer, 500 µL of High Salt Wash Buffer, and two final washes with 500 µL of Wash Buffer.

The samples were then denatured by adding 7.5 µL of 4× LDS buffer (ThermoFisher, NP0008), 3 µL of 1 M DTT, and 20 µL of Wash Buffer, followed by heating at 70°C for 10 minutes. Proteins were separated via NuPAGE gel electrophoresis, transferred to nitrocellulose membranes (Amersham Protran, 0.45 µm), and the region corresponding to 100–250 kDa was excised. Protein digestion was performed using 150 µL of Proteinase K SDS mix at 37°C for 20 minutes, followed by an additional incubation at 50°C for another 20 minutes.

RNA was extracted by phenol:chloroform:isoamyl alcohol (PCA) precipitation and resuspended in 9 µL of nuclease-free water. Reverse transcription was initiated by adding 1 µL of 10 mM dNTPs and 0.5 µL of RTP primer to the RNA, heating the mixture at 65°C for 3 minutes, and then cooling it on ice. A 10 µL RT mix was then added, and the reaction was incubated at 55°C for 20 minutes.

To remove unused primers and nucleotides, 2.5 µL of ExoSAP-IT (ThermoFisher, 78201.1) was added and incubated at 37°C for 15 minutes. The mixture was then treated with 1 µL of 0.5 M EDTA and 3 µL of 1 M NaOH at 70°C for 10 minutes, followed by neutralization with 3 µL of 1 M HCl.

The resulting cDNA was purified using 5 µL of Silane beads (Invitrogen, 37002D) in a mixture of 90 µL RLTW buffer and 108 µL of 100% ethanol. Beads were incubated at room temperature for 10 minutes and then washed twice with 300 µL of 80% ethanol, followed by a final wash with 150 µL of 80% ethanol. After air-drying for 5 minutes, the cDNA was prepared for 5′ adapter ligation.

For this, cDNA was denatured in 1.45 µL of TT Elution Buffer, 0.5 µL of RA5 DNA adapter (100 µM), and 0.8 µL of DMSO at 70°C for 2 minutes. Then, 7.55 µL of 5′ adapter ligation mix was added, and ligation proceeded overnight at room temperature with gentle rotation.

Following ligation, a second round of silane bead purification was performed using 2.5 µL beads, 45 µL of RLTW buffer, and 45 µL of ethanol. After ethanol washes as above, the cDNA was eluted in 22 µL of Bead Elution Buffer.

Finally, cDNA libraries were amplified by PCR using RPI1 and RPIX primers for 12 cycles. The amplified products were size-selected on 8% polyacrylamide gels to enrich for fragments between 180 and 400 bp. Sequencing was carried out on the Illumina NovaSeq platform.
